# Metagenomic profiling of ammonia- and methane-oxidizing microorganisms in a Dutch drinking water treatment plant

**DOI:** 10.1101/2020.05.19.103440

**Authors:** Lianna Poghosyan, Hanna Koch, Jeroen Frank, Maartje A.H.J. van Kessel, Geert Cremers, Theo van Alen, Mike S.M. Jetten, Huub J.M. Op den Camp, Sebastian Lücker

## Abstract

Elevated concentrations of ammonium and methane in groundwater can cause severe problems during drinking water production. To avoid their accumulation, raw water in the Netherlands, and many other countries, is purified by sand filtration. These drinking water filtration systems select for microbial communities that mediate the biodegradation of organic and inorganic compounds. In this study, the active layers and wall biofilm of a Dutch drinking water treatment plant (DWTP) were sampled at different locations along the filtration units of the plant over three years. We used high-throughput sequencing in combination with differential coverage and sequence composition-based binning to recover 56 near-complete metagenome-assembled genomes (MAGs) with an estimated completion of ≥70% and with ≤10% redundancy. These MAGs were used to characterize the microbial communities involved in the conversion of ammonia and methane. The methanotrophic microbial communities colonizing the wall biofilm (WB) and the granular material of the primary rapid sand filter (P-RSF) were dominated by members of the *Methylococcaceae* and *Methylophilaceae.* The abundance of these bacteria drastically decreased in the secondary rapid sand filter (S-RSF) samples. In all samples, complete ammonia-oxidizing (comammox) *Nitrospira* were the most abundant nitrifying guild. Clade A comammox *Nitrospira* dominated the P-RSF, while clade B was most abundant in WB and S-RSF, where ammonium concentrations were much lower. In conclusion, the knowledge obtained in this study contributes to understanding the role of microorganisms in the removal of carbon and nitrogen compounds during drinking water production. We furthermore found that drinking water treatment plants represent valuable model systems to study microbial community function and interaction.

**Highlights:**

- Microbial distribution was mainly influenced by sampling location within the DWTP
- Clade A comammox *Nitrospira* were the dominant nitrifiers in the primary sand filter
- Clade B was most abundant in samples from wall biofilm and the secondary filter
- A novel *Methylophilaceae*-affiliated methanotroph dominated the primary sand filter

## Introduction

About 97% of all available water on earth is saline. The remaining 3% is freshwater, of which more than two-thirds is frozen in ice sheets. Thus, only a small fraction of the global freshwater exists as ground and surface water that is available for drinking water production. According to the European Commission (EC, 2016), about 50% of drinking water in Europe is produced from groundwater and 37% from surface water.

Groundwater has a relatively constant composition and may contain high concentrations of (Fe^2+^; 0.9-7.8 mg/L), manganese (Mn^2+^; 0-0.56 mg/L), ammonium (NH_4_^+^; 0.1-0.5 mg/L), and some organic compounds such as methane (CH_4_; 0-37 mg/L) (Albers et al., 2015; Li and Carlson, 2014; Osborn et al., 2011). Elevated concentrations of these compounds in groundwater can cause severe problems during drinking water production and distribution (Okoniewska et al., 2007; Rittmann et al., 2012; Sharma et al., 2005). Biofiltration (e.g., rapid (RSF) or slow (SSF) sand filtration, granular activated carbon filters) are widely applied methods for the removal of the above-mentioned compounds. Biofilters harbor complex microbial communities that are introduced via the source water (Yang et al., 2016) and are shaped by the configuration of the treatment process (Li et al., 2017; Pinto et al., 2012). In the filtration units microbial growth is stimulated on filter material, mediating the biodegradation of organic and inorganic compounds (Proctor and Hammes, 2015). Gases such as methane, hydrogen sulfide, carbon dioxide, and other volatile compounds are removed from the groundwater through gas exchange systems (Trussell et al., 2012). The increased dissolved oxygen in the water caused by this mechanical aeration step serves as an electron acceptor in microbially mediated oxidative reactions, which may ensure the near-complete nutrient removal in the biologically active layer of the sand filters.

One of the main groundwater contaminants is ammonium. Excess ammonium in raw water is often associated with microbiological, chemical and sanitary problems in drinking water distribution systems, such as excessive biofilm growth, pH decrease, pipe corrosion, and elevated nitrite and nitrate levels (Beech and Sunner, 2004; Camper, 2004; Rittmann et al., 2012). In engineered systems, such as drinking water treatment plants (DWTP), ammonium removal is achieved by the activity of nitrifying microorganisms that oxidize ammonia to nitrate via a series of intermediates. While canonical nitrifying guilds perform ammonia- and nitrite-oxidation in a tight interplay, complete ammonia-oxidizing (comammox) bacteria of the genus *Nitrospira* possess all proteins necessary to perform nitrification on their own (Daims et al., 2015; van Kessel et al., 2015). Nitrifying microbial communities of rapid sand filters have been studied before and seem to be represented by different groups of nitrifiers (Albers et al., 2015; Fowler et al., 2018; Gülay et al., 2016; Oh et al., 2018; Palomo et al., 2016; Pinto et al., 2015; van der Wielen et al., 2009). In these systems, nitrification can be limited by the availability of vital nutrients for this process, such as phosphate and copper. This can cause incomplete nitrification (de Vet et al., 2012; Wagner et al., 2016), leading to incomplete ammonium removal and/or nitrite accumulation (Wilczak et al., 1996), and microbial after-growth in the distribution network (Rittmann et al., 2012).

Methane is a colorless and odorless gas that usually does not present a health risk in drinking water. However, methane gas is highly flammable and can be explosive at elevated concentrations, and it also can serve as substrate for growth of microorganisms in distribution systems. Most of the methane is removed by the mechanical aeration step and remaining amounts are oxidized by aerobic methanotrophs within the sand beds of the filter units. Correspondingly, a number of studies reported methane-oxidizing bacteria colonizing the granular material of the sand filters (Albers et al., 2015; Gülay et al., 2016; Palomo et al., 2016). However, in contrast to nitrifying microbial communities, methanotrophs in these engineered systems are traditionally less well studied. Methane is a potent greenhouse gas, with a global warming potential of 34 CO_2_ equivalents over 100 years (Stocker et al., 2013) and the removal of most methane from the drinking water via aeration causes methane emissions to the atmosphere, contributing to global warming and climate change (Maksimavičius and Roslev, 2020).

Like in many European countries, groundwater is the primary source (65%) for drinking water production in the Netherlands (Dutch Drinking Water Statistics 2017). In this study, we used genome-resolved metagenomics and gene-centric approaches to analyze microbial communities in a Dutch DWTP, with a special focus on ammonia- and methane-oxidizing microorganisms. The groundwater entering this DWTP contains elevated concentrations of organic (CH_4_, 5.2 mg/L) and inorganic (NH_4_^+^, 0.66 mg/L; Fe^2+^, 8.4 mg/L; Mn^2+^, 0.18 mg/L) nutrients, which are removed in two sequential rapid sand filters. Thus, this DWTP represents an interesting model system to study microbial communities involved in the conversion and removal of these compounds. Metagenomic analysis of samples taken at different stages within this DWTP revealed the key microorganisms involved in ammonia- and methane-oxidation, including novel methanotrophic bacteria from the *Methylophilaceae* family, which were assumed previously to comprise only methylotrophic bacteria.

## Materials and Methods

### Sample collection and trace element analysis

Samples were obtained from the pumping station Breehei, a drinking water treatment facility located in Venray, the Netherlands (51°28′54.6″N; 5°59′10.2″E), operated by NV Waterleiding Maatschappij Limburg. Drinking water is produced from groundwater. Samples from the active layers of primary and secondary rapid sand filters (P-RSF and S-RSF) were collected in 50 ml sterile falcon tubes in June 2016 and September 2018. Samples from the biofilm formed on the walls of the primary sand filter (WB) were taken in June 2016, May 2017 and September 2018 (Figure 1). All samples were transferred to the laboratory within 4 h, and were stored at 4°C for further analysis. Water quality parameters were determined by Aqualab Zuid (https://www.aqualab.nl) over the period 2000-2016 (Table 1).

**Figure 1.**
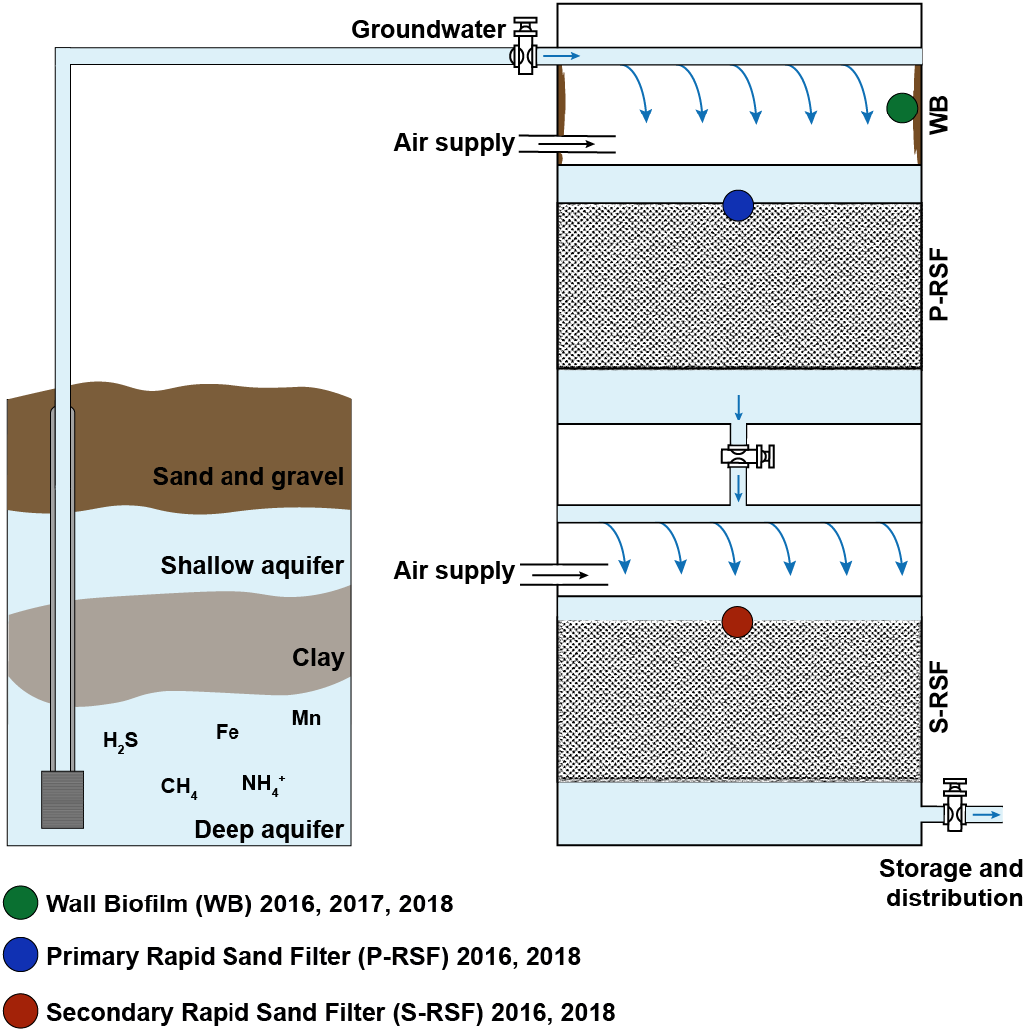
Schematic illustration of DWTP Breehei. Sampling locations are indicated by colored dots, sampling timepoints (in years only) for each location are indicated.

**Table 1.** Average water quality parameters of the incoming groundwater and at different stages along the treatment train.

| Unit (mg L<sup>-1</sup>) | Ground-water | After aeration | P-RSF effluent | S-RSF effluent | Drinking water | WML standard<sup>a</sup> |
| --- | --- | --- | --- | --- | --- | --- |
| **Methane (CH<sub>4</sub>)** | 5.2 | 1.2 | 0.012 | 0.012 | 0.012 | 0.5 |
| **Ammonium (NH<sub>4</sub><sup>+</sup>)** | 0.66 | 0.65 | 0.0145 | 0.039 | 0.039 | 0.1 |
| **Nitrite (NO<sub>2</sub><sup>-</sup>)** | 0.0046 | * | 0.063 | 0.0082 | 0.0041 | 0.05 |
| **Nitrate (NO<sub>3</sub><sup>-</sup>)** | 0.13 | * | * | * | 1.64 | 40 |
| **Oxygen (O<sub>2</sub>)** | 0.4 | 9.1 | 5.0 | 10.2 | 10.2 | - |
| **Carbon dioxide (CO<sub>2</sub>)** | 42.38 | * | 18 | 6.16 | 6.95 | - |
| **Iron (Fe)** | 8.4 | * | 0.11 | 0.053 | 0.043 | 0.1 |
| **Manganese (Mn)** | 0.18 | * | 0.022 | 0.0079 | 0.0052 | 0.025 |

\* Not applicable or were not analyzed
- No standards
<sup>a</sup>Defined by Waterleiding Maatschappij Limburg

### Methane uptake

Methane-oxidizing capacity was determined for the P-RSF and WB samples collected in 2016. For the P-RSF, 2.5, 5, 10, and 20 g of sand material were mixed with 20 ml top water. After settling of the sand, overlaying water samples (20 ml) were transferred into 120 ml serum bottles with ±14.5 mg/L CH_4_ in the headspace and incubated at room temperature in a shaking incubator (200 rpm). For the WB sample, the incubations were conducted in triplicates using 1 and 2.5 g of biomass. CH_4_ concentrations were determined by sampling 300 μL headspace, which were injected in triplicates into a HP 5890 gas chromatograph (Hewlett Packard, Palo Alto, CA).

### DNA extractions

DNA from samples for Illumina sequencing was extracted using two different methods to obtain differential abundance information. DNA from 2016 samples was extracted with DNeasy Blood & Tissue Kit (Qiagen Ltd., West Sussex, United Kingdom) and PowerSoil DNA Isolation Kit (MO BIO Laboratories, Carlsbad, CA, USA). Samples (0.5 g) were mechanically disrupted using a TissueLyser (Qiagen) for 2 × 30 seconds at 30 Hz, followed by DNA extractions according to the manufacturer’s instructions. For the extraction of DNA from samples collected in 2017 and 2018, the DNeasy Blood & Tissue kit was replaced by ammonium acetate (Kowalchuk et al., 2004) and CTAB (Zhou et al., 1996) based extraction methods, respectively.

For long read Nanopore sequencing, DNA was extracted from P-RSF samples collected in 2016 and 2018 using the CTAB-based extraction method. To avoid shearing of genomic DNA, all bead-beating and vortexing steps were replaced by carefully inverting the tubes several times, and all the pipetting steps were performed using cut-off pipette tips. Genomic DNA was purified twice by phenol:chloroform:isoamyl alcohol (25:24:1) phase extraction (Zhou et al., 1996). Extracted DNA was resuspended in nuclease-free water and stored at 4°C.

### Illumina library preparation and sequencing

For Illumina library preparation, the Nextera XT kit (Illumina, San Diego, CA, USA) was used according to the manufacturer’s instructions. Enzymatic tagmentation was performed using 1 ng of DNA per sample, followed by incorporation of the indexed adapters and library amplification. After subsequent purification using AMPure XP beads (Beckman Coulter, Indianapolis, IN, USA), libraries were checked for quality and size distribution using the 2100 Bioanalyzer with the High Sensitivity DNA kit (Agilent, Santa Clara, CA, USA). Library quantitation was performed by Qubit using the Qubit dsDNA HS Assay Kit (Thermo Fisher Scientific, Waltham, MA, USA). After dilution to 4 nM final concentration, the libraries were pooled, denatured, and sequenced on an Illumina Miseq. Paired-end sequencing of 2 × 300 base pairs was performed using the Illumina MiSeq Reagent Kit v3 according to the manufacturer’s protocol.

### Nanopore library preparation and sequencing

For Nanopore library preparation, 1 - 1.5 μg of DNA, measured by Qubit with the Qubit dsDNA HS Assay Kit (Thermo Fisher Scientific, Waltham, MA, USA), was used. The input DNA was quality-checked by agarose gel electrophoresis to contain only high molecular DNA and show no degradation. For sequencing, DNA Library construction was performed using the Ligation Sequencing Kit 1D (SQK-LSK108) in combination with the Native Barcoding Expansion Kit (EXP-NBD103 or EXP-NBD104) according to the manufacturer’s protocol (Oxford Nanopore Technologies, Oxford, UK). Fragments were end-repaired using the NEBNext^®^ FFPE DNA Repair Mix (New England Biolabs, Ipswich, MA, USA), with subsequent fragment purification using AMPure XP beads (Beckman Coulter Life Sciences, Indianapolis, IN, USA). End repair and dA-tailing was done using the NEBNext^®^ Ultra™ II End Repair/dA-Tailing Module (New England Biolabs) followed by a cleanup of the fragments using AMPure XP beads. Selected barcodes for each sample were ligated using the Blunt/TA Ligase Master Mix (New England Biolabs) and the resulting fragments were purified using AMPure XP beads. The DNA concentration of all libraries was measured by Qubit using the dsDNA HS Assay Kit and pooled to a maximum of 700 ng DNA. Subsequently, adapters were ligated using the NEBNext^®^ Quick Ligation Module (New England Biolabs). After purification using AMPure XP beads, the pooled libraries were quantified again using Qubit. The libraries were loaded on a Flow Cell (R9.4.1) and run on a MinION device (Oxford Nanopore Technologies, Oxford, UK), according to the manufacturer’s instructions. Base calling after sequencing was done using Albacore v2.1.10, (Oxford Nanopore Technologies) for the 2016 and guppy_basecaller in combination with guppy_barcoder (Oxford Nanopore Technologies, Limited Version 2.3.7+e041753) for the 2018 sample.

### Assembly

Raw Illumina sequence data were processed for quality-trimming, adapter removal, and contamination-filtering, using BBDUK (BBTOOLS v37.17; http://jgi.doe.gov/data-and-tools/bbtools/bb-tools-user-guide/bbduk-guide/). All sequencing reads from the same sampling location were co-assembled using metaSPAdes v3.10.1 (Nurk et al., 2017) with the following parameters: k-mer sizes 21, 33, 55, 77, 99 and 127, minimum contig length 1500 bp. Raw Nanopore reads were quality trimmed using Filtlong v0.2.0 (https://github.com/rrwick/Filtlong) with minimum read length 1000 bp and error rate <20%. Porechop v0.2.4 (https://github.com/rrwick/Porechop) was used to remove adapters and split chimeric reads with default settings. Trimmed reads were assembled using Canu v1.8 (Koren et al., 2017) with minimum read length 1000, corrected read error rate 0.105, and genome size 5m. To correct error rates, the trimmed Nanopore reads were mapped to the assembly using Minimap2 v2.16-r922 (Li, 2018), which then was polished with Racon v1.3.1 (Vaser et al., 2017). Subsequently, Racon v1.3.1 was used to further polish the assembly twice with Illumina reads obtained from the 2018 P-RSF sample.

### Metagenome binning

Differential coverage information was determined by separately mapping the sequencing reads from each sample and DNA extraction method against the obtained co-assemblies, using Burrows-Wheeler Aligner v0.7.15 (BWA) (Li and Durbin, 2010) and employing the “mem” algorithm. For Canu assembly, only the Illumina reads from P-RSF 2018 samples were mapped. The generated sequence alignment map (SAM) files were converted to binary format (BAM) using SAMtools (Li et al., 2009). Metagenome binning was performed using anvi′o v5.3 (Eren et al., 2015). Anvi′o′s Snakemake-based (Koster and Rahmann, 2012) contigs workflow was used to generate contig databases from each Illumina and Nanopore assembly. Shortly, anvi′o employs Prodigal v2.6.3 (Hyatt, 2010) to identify open reading frames (ORFs) and HMMER v3.2 (Eddy, 2011) to identify archaeal (Rinke et al., 2013) and bacterial (Campbell et al., 2013) single-copy core genes. The Cluster of Orthologous Groups of proteins (COG) database (Tatusov et al., 1997) together with Centrifuge v1.0.3-beta (Kim et al., 2016) was used to annotate genes in the contig databases. Each BAM file was profiled to obtain differential coverage and statistical information based on mapping results and to generate merged profile databases. The contigs were then automatically clustered with CONCOCT (Alneberg et al., 2014), followed by manual binning and bin refinement using the anvi′o interactive interface (Delmont and Eren, 2016; Eren et al., 2015). Completeness and contamination (referred to as redundancy in this study) of bins was assessed by CheckM v1.01.11 (Parks et al., 2015), which uses pplacer v1.1 alpha 19 (Matsen et al., 2010) to identify and quantify single-copy marker genes. Based on the suggested standards (Bowers et al., 2017), the bins were defined as high-quality (>90% complete and <5% redundancy, complete small subunit rRNA operon, ≥18 tRNAs) and medium-quality (≥70% complete and <10% redundancy) metagenome-assembled genomes (MAGs).

### Dereplication and taxonomic classification of MAGs

MAGs were dereplicated using dRep v2.2.3 (Olm et al., 2017) at 99% average nucleotide identity (ANI) for clustering. Within each cluster, the best MAG was selected based on completeness (≥70%), redundancy (<10%), N50 of contigs, and fragmentation. GTDB-Tk v0.3.2 (Parks et al., 2018) was used for taxonomic assignment of the final MAGs. Phyla are named according to the recently suggested nomenclature (Whitman et al., 2018) using standardized phylum suffix -ota.

### Abundance estimation of MAGs

To calculate the relative abundance of the dereplicated MAGs in each sample, reads from all samples were individually mapped to each co-assembly using BWA v0.7.15 (Li and Durbin, 2010) as described above. The coverage of each MAG was calculated using CheckM v1.01.11 (minimum alignment length 0.95) (Parks et al., 2015) and was normalized by multiplying this coverage with a normalization factor (sequencing depth of the largest sample divided by the sequencing depth of each individual sample). The distribution of MAGs was calculated as percentages by dividing a MAG’s coverage in each sample by the total coverage of the respective MAG in all samples.

### Functional analysis

For the gene-centric approach, co-assembly across all samples was performed using MEGAHIT v1.1.1-2 (Li et al., 2015). Open reading frames (ORFs) in the MAGs obtained above and the new co-assembly were predicted using Prodigal v2.6.3 (Hyatt, 2010), which was set to include partial ORFs. Custom-build hidden Markov models (HMMs) (Eddy, 2011) of specific marker proteins were used (Supplementary material and methods) to annotate all ORFs using hmmsearch (HMMER v3.1b2; http://hmmer.org). The HMM for RNA polymerase subunit beta (RpoB) was downloaded from FunGene (Fish et al., 2013). Remaining ORFs in the MAGs were annotated using Prokka v1.12-beta (Seemann, 2014). The annotations of all genes discussed in this study were confirmed by BLAST against the TrEMBL, Swiss-Prot and NCBI nr databases. Subcellular localization of the proteins was predicted by SignalP 5.0 (Armenteros et al., 2019) and TMHMM 2.0 (Krogh et al., 2001).

Functional gene-based abundances of ammonia- and methane-oxidizing microorganisms were estimated using competitive metagenomic read recruitment to ensure unique mapping. For this, reads from each metagenomic sample were mapped using bowtie2 v2.3.1 (Langmead and Salzberg, 2012) in ‘-very-sensitive’ mode against extracted partial and complete sequences of *rpoB* and the ammonia as well as the particulate and soluble methane monooxygenase subunit A/alpha genes (*amoA*, *pmoA* and *mmoX*, respectively). SAMtools flagstat v1.6 (Li et al., 2009) was used to obtain the number of mapped reads. Reads per kilo base per million mapped reads (RPKM)-values were used to correct for differences in sequencing depth and gene length. To estimate the relative abundance of microorganisms encoding ammonia and methane monooxygenases in each sample, the normalized read counts were calculated as fraction of the normalized read counts of the identified *rpoB* genes.

### Phylogenomic and phylogenetic analyses

The up-to-date bacterial core gene (UBCG) pipeline (Na et al., 2018) with default parameters was used to extract and concatenate bacterial core gene sets. To infer the phylogeny of the archaeal MAGs, the anvi’o phylogenomic workflow (http://merenlab.org/2017/06/07/phylogenomics) was used to individually align and concatenate 162 archaeal single-copy genes. Maximum-likelihood trees were calculated using RAxML version 8.2.10 (Stamatakis, 2014) on the CIPRES science gateway (Miller et al., 2010). For details, see Supplementary Methods. Amino acid sequences of type II DMSO reductase-family enzymes, and of ammonia and methane monooxygenases were aligned using ARB v5.5 (Ludwig et al., 2004). The maximum-likelihood trees were calculated using RAxML HPC-HYBRID v.8.2.12 on the CIPRES and IQ-tree webserver (Trifinopoulos et al., 2016) as described in Supplementary Methods. All phylogenetic trees were visualized in iTOL (Letunic and Bork, 2016).

### Data visualization

Manuscript figures were generated using ggplot2 (Wickham, 2016), Rstudio (Racine, 2012) and the anvi’o interactive interface (http://merenlab.org/2016/02/27/the-anvio-interactive-interface).

## Results

### DWTP performance

The produced water quality analyses (Table 1) showed that methane, ammonium, nitrite and nitrate were well below the required quality standards. Most of the methane in the raw water was removed during the aeration step and the remainder was oxidized in the P-RSF. Pronounced ammonia oxidation in P-RSF resulted in nitrite accumulation in the effluent of this filter, which was subsequently removed in the S-RSF. Similarly, the Fe^2+^ and Mn^2+^ removal efficiency of the system was very high (>99%). In addition, we determined methane uptake rates in the P-RSF and WB samples. Complete methane oxidation was achieved in all samples within 5 to 8 days of incubation, except for the control containing only raw water (Figure S1). Methane consumption in P-RSF samples increased with an increasing amount of sand material, whereas no significant difference in methane uptake was detected between the incubations with 1 and 2.5 g of WB biomass (Figure S1). The identical oxidation rates in the incubations with WB biomass seem counterintuitive, but may be caused by sample inhomogeneity and uneven distribution of methanotrophic bacteria within the biofilm.

### Recovery of metagenome-assembled genomes

Over the period 2016 to 2018 a total of 7 samples from DWTP Breehei were collected (Figure 1) and sequenced. This resulted in a total of 125 million paired-end Illumina sequencing reads. For each sample location more than 70% of the respective reads could be co-assembled into ~413, 184, and 249 Mb sequencing data for WB, P-RSF, and S-RSF, respectively (Table S1). In addition, P-RSF samples were also sequenced using the Oxford Nanopore long-read platform to improve the assembly of the most abundant microorganisms. Overall, binning of 4 individual metagenome assemblies based on sequence composition and differential coverage patterns resulted in 78 near-complete Illumina and 7 Nanopore MAGs (Table S2). All MAGs obtained from Illumina co-assemblies as well as the Nanopore assembly were dereplicated at strain level (99% ANI), which yielded 50 medium and 6 high-quality (MIMAG standards; Bowers et al., 2017) non-redundant MAGs (Table 2, Table S2) that were used for downstream analyses. Given the high number of single nucleotide variants observed during bin refinement using the anvi′o interactive interface, 15 MAGs were categorized as population-level genomes (Table 2). All MAGs were classified at the lowest possible taxonomic level using GTDB-tk (Parks et al., 2018), indicating affiliation with 1 archaeal and 12 different bacterial phyla (Figure 2, Table 2).

**Figure 2.**
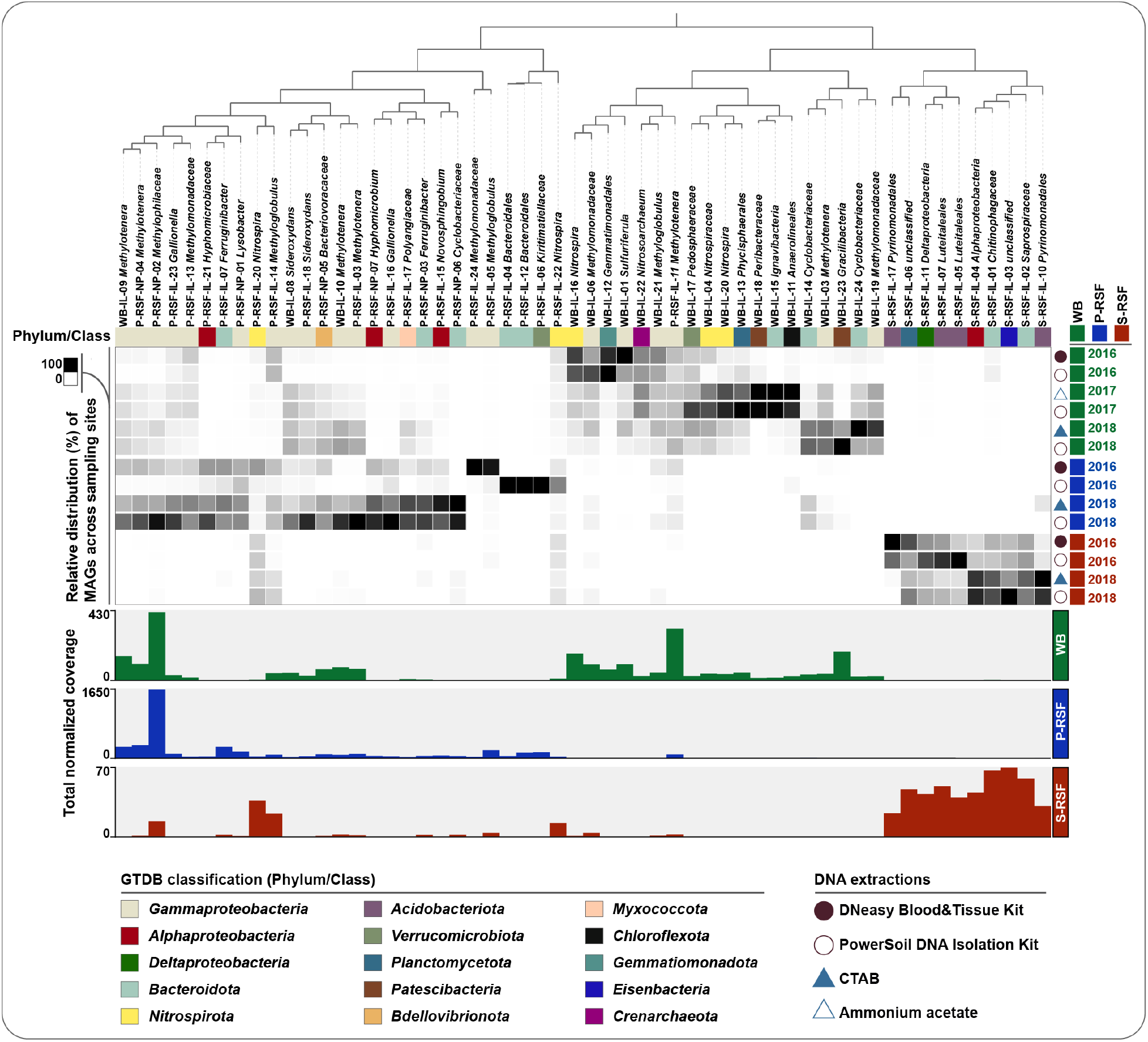
Overview of the distribution, coverage and classification of 56 dereplicated medium- and high-quality MAGs recovered from the DWTP Breehei metagenomes. MAGs are organized based on their relative distribution in the DWTP sample locations using Euclidean distance and Ward linkage as implemented in the anvi′o interactive interface. The phylogenetic affiliation of the MAGs based on GTDB classification is indicated by colored boxes. The heatmap indicates the percent coverage of each MAG in a given metagenome. The bar plots show the total normalized coverage of each MAG per sampling site.

**Table 2.**
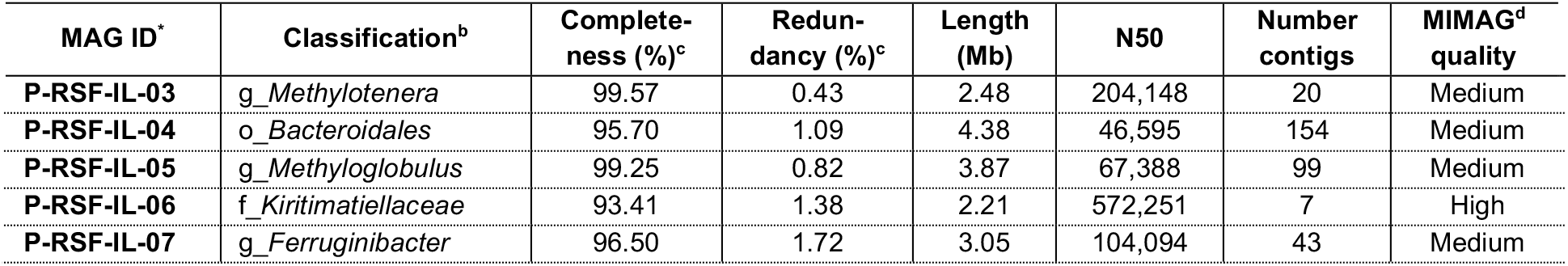
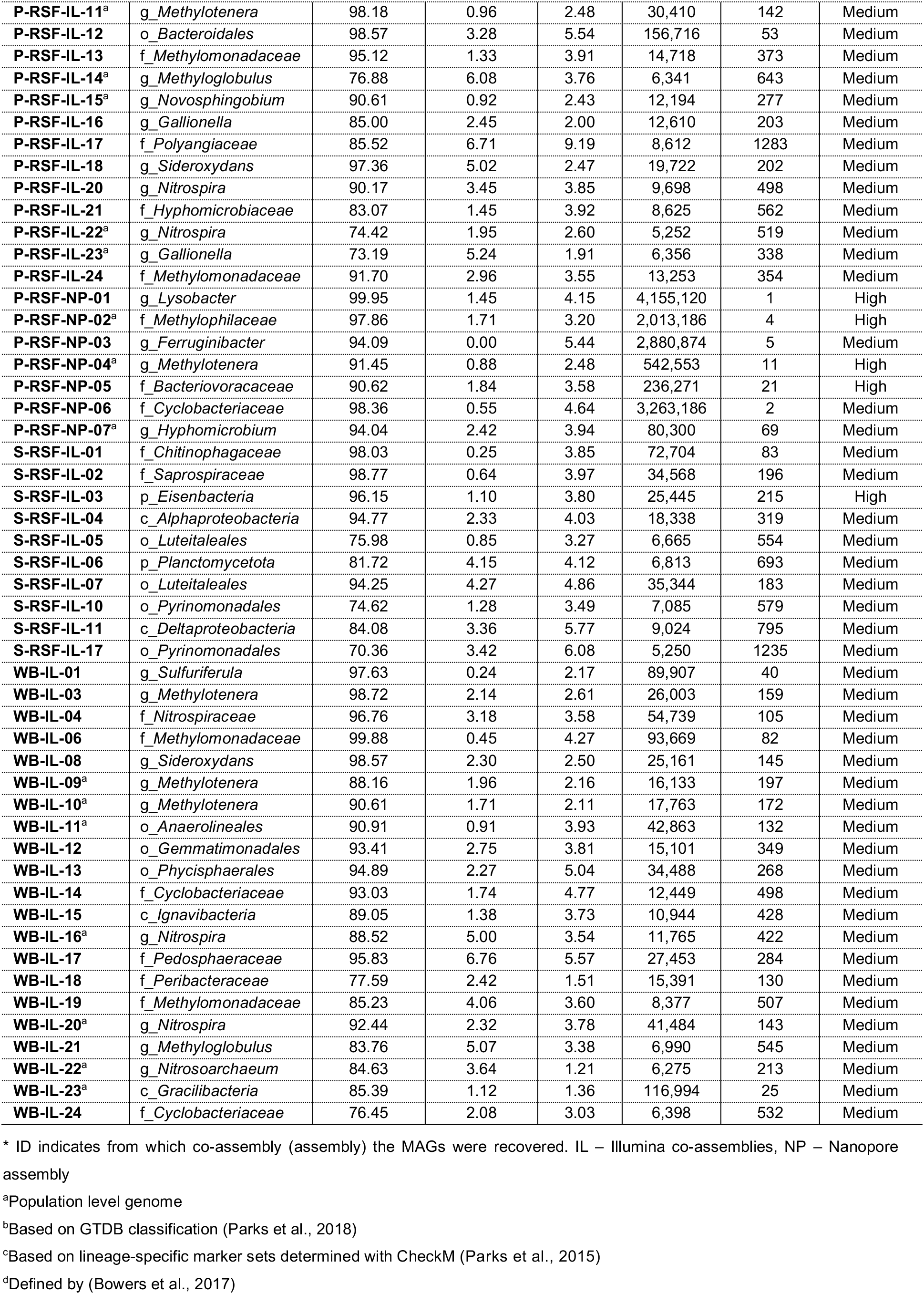
General characteristics of MAGs recovered from DWTP Breehei metagenomes.

### Distribution and taxonomic composition of the DWTP microbiome

The influence of sampling location, sampling time, and DNA extraction method on the distribution of recovered MAGs was analyzed by hierarchical clustering using Euclidean distance metrics with a Ward linkage algorithm, allowing grouping of the MAGs based on their occurrence patterns in the different samples (Figure 2). Overall, the choice of a DNA extraction method had no pronounced influence on the distribution of MAGs across samples, except for the samples from P-RSF in 2016. In this specific sample, the normalized coverage values indicated a strong extraction bias for DNA extracted using the Power soil (P-RSF16_PS) compared to the Blood and Tissue kit (P-RSF16_BT; Table S2). Notably, this bias was not observed for the 2016 S-RSF samples extracted with the same kits, or for any other DNA extraction performed using the Power soil kit. It thus is difficult to conceive that the extraction method was affecting specific members only in the 2016 P-RSF microbial community, and the reason for the observed bias remains unclear. Overall, the sampling location had the most substantial effect on microbial diversity and abundance. While some MAGs recovered from WB and P-RSF were also present in S-RSF, the overall community of the S-RSF community clearly differed from the other sampling locations (Figure 2). The microbial communities of WB and P-RSF were generally more similar, but sampling location and time influenced the relative abundance of MAGs across all samples.

To gain insights into the overall microbial community structure and diversity of the DWTP Breehei, 16S rRNA gene sequences were retrieved directly from the metagenomic assemblies. Both full-length and partial 16S rRNA gene sequences were used for further analyses, since only 21 to 32% of 16S rRNA reads were assembled into full-length sequences (Figure S2). Subsequently, microbial community composition was analyzed at the phylum and family levels (Figure S3). Although changes in abundance were observed, phylum level classification revealed no differences in microbial community composition between the sampling locations (Figure S3-A). At the family level, the S-RSF microbial community was clearly distinct from the WB and P-RSF community (Figure S3-B), corroborating the MAG co-occurrence pattern-based observations when samples were organized using Euclidian distance and Ward ordination (Figure 2).

16S rRNA gene-based taxonomic profiling identified 14 phyla with relative abundances ≥1% (Figure S3A). *Gammaproteobacteria* (17-67%), *Bacteroidota* (5-17%), *Acidobacteria* (2-15%), *Alphaproteobacteria* (4-12%), *Planctomycetota* (2-10%) *“Ca.* Patescibacteria” (CPR; 1-7%), and *Nitrospirota* (2-6%) were the most dominant phyla identified in all samples (Figure S3-A). In most cases, this 16S rRNA gene-based analysis corresponded well to the taxonomic affiliation of the recovered MAGs. Additionally, one MAG belonging to the candidatus phylum “*Ca.* Eisenbacteria” and one *Crenarchaeota* (*Thaumarchaeota*) MAG were obtained from WB and S-RSF (Figure 1), but were of low abundance in the 16S rRNA datasets.

### Genome functional profiling

Iron, manganese, reduced sulfur species, ammonium and methane are removed during the drinking water treatment process. However, our understanding of the microbial and geochemical processes contributing to removal of these compounds during sand filtration is still limited. Although iron (Fe^2+^) oxidation pathways in bacteria are not fully understood, it is known that certain autotrophic bacteria are responsible for this process. Four *Gallionellaceae* MAGs (2 *Gallionella,* 2 *Sideroxydans*) were present only in WB and P-RSF metagenomes (Figure 2), indicating that complete iron removal may already occur in P-RSF. The potential to use reduced sulfur compounds as electron donor was encoded in *Hyphomicrobiaceae* (P-RSF-IL-21) and *Sulfuriferula* (WB-IL-01) MAGs (Table S3). To learn more about the microorganisms driving the removal of methane and ammonium in this rapid sand filtration system, a genome-resolved metagenomics approach was applied to recover high-quality genomes of the key ammonia- and methane-oxidizing microorganisms. Subsequently, all recovered MAGs were screened for functional marker proteins involved in nitrification, and methane and other one-carbon (C1) compound oxidation (Figure 3, Table S3) using custom-build HMM models.

**Figure 3.**
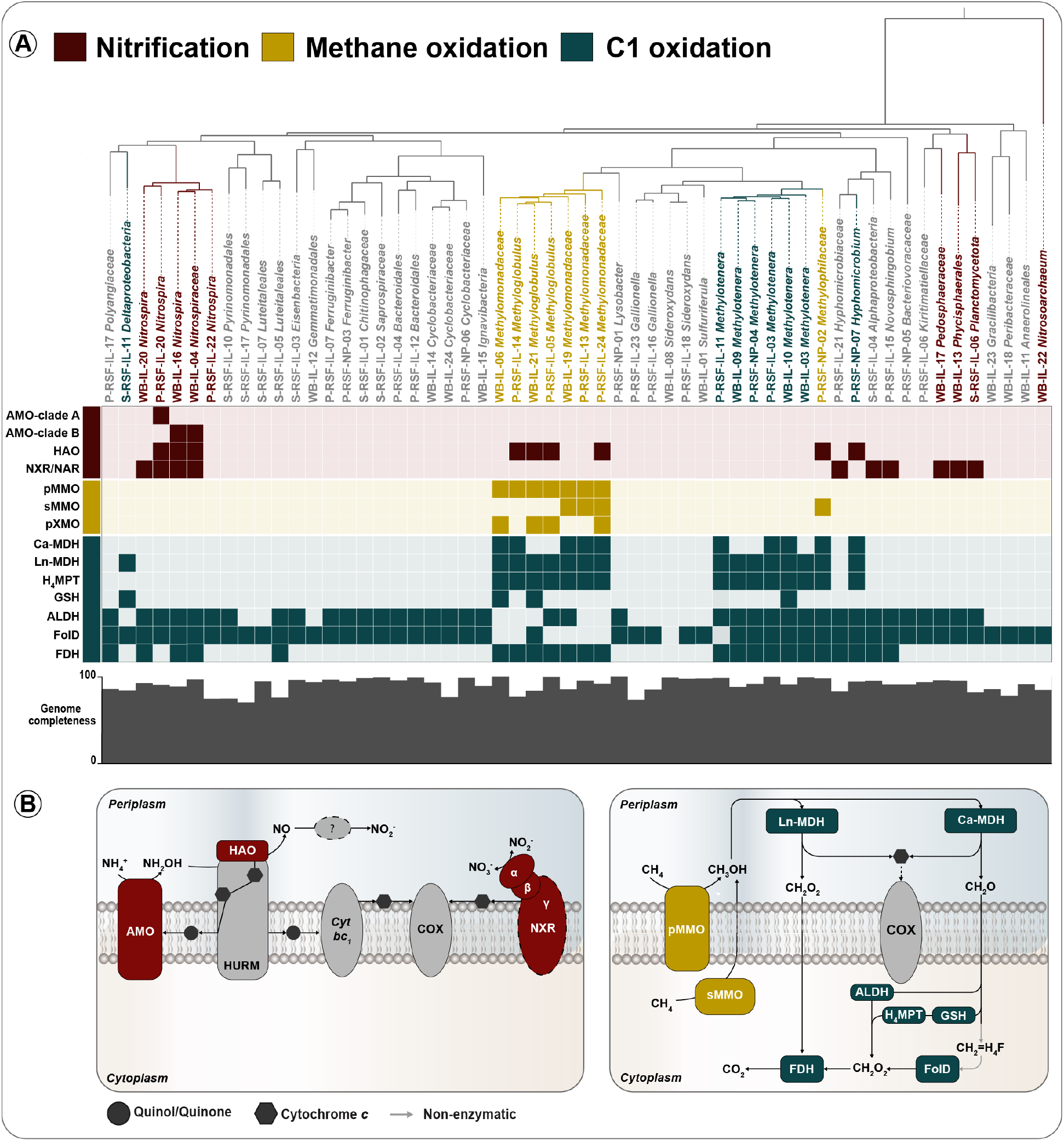
Metabolic potential of the 56 MAGs recovered from DWTP Breehei **A.** Phylogenetic tree based on the concatenated alignment of 49 ribosomal proteins (Table S3) using the anvi′o phylogenomics workflow. Presence or absence of genes for ammonia, nitrite, methane, and C1 utilization are indicated by filled or shaded colored boxes, respectively. Grey bars represent estimated genome completeness. The figure was generated using the anvi′o interactive interface. **B.** Schematic pathway models for complete nitrification, and methane and C1 oxidation, based on enzyme complexes identified in the metagenome. ALDH, aldehyde dehydrogenase; AMO, ammonia monooxygenase; COX, cytochrome *c* oxidase; FDH, formate dehydrogenase; FolD, methylenetetrahydrofolate dehydrogenase/cyclohydrolase; GSH, glutathione-linked formaldehyde oxidation; HAO, hydroxylamine dehydrogenase; H4MPT, tetrahydromethanopterin; HURM, hydroxylamine-ubiquinone redox module; pMMO, particulate methane monooxygenase; sMMO, soluble methane monooxygenase; MDH, lanthanide and calcium-dependent methanol dehydrogenases; NAR, nitrate reductase; NXR, nitrite oxidoreductase.

### Ammonia and nitrite oxidation

16S rRNA gene sequence analysis revealed the presence of nitrifying bacteria affiliated with the families *Nitrospiraceae* and *Nitrosomonadaceae* in the DWTP Breehei. *Nitrospiraceae* dominated the nitrifying microbial community in all samples, but their abundance patterns differed along the different sampling locations, with the lowest abundance in P-RSF. In contrast, canonical ammonia-oxidizing bacteria (AOB) affiliated with the *Nitrosomonaceae* showed very low abundance in all samples (Figure S3). Metagenomic consensus binning allowed the recovery of five *Nitrospira* MAGs from WB and P-RSF and one MAG of an ammonia-oxidizing archaeum (AOA) affiliated with the genus *Nitrosoarchaeum*. Despite the detection of *Nitrosomonadaceae* in WB and S-RSF samples in 16S rRNA-based analyses, no high or medium-quality metagenomic bin of this taxonomic group was recovered.

All MAGs were screened for key genes of autotrophic nitrification, including the gene sets encoding ammonia monooxygenase (AMO) and nitrite oxidoreductase (NXR). The AMO complex catalyzes the oxidation of ammonia to hydroxylamine (Figure 3B) and belongs to the family of copper-containing membrane monooxygenases (CuMMOs or XMO) (Khadka et al., 2018). Based on the phylogeny of AMO subunit A (AmoA), comammox *Nitrospira* form two monophyletic clades referred to as clade A and B (Daims et al., 2015). AmoA is often used as a functional and phylogenetic marker for ammonia-oxidizing microorganisms (Junier et al., 2008; Pester et al., 2012; Pjevac et al., 2017) and thus AmoA was used to examine the full ammonia oxidation potential in the DWTP Breehei. Phylogenetic analysis placed our metagenome-derived AmoA sequences into five divergent groups (Figure 4A).

**Figure 4.**
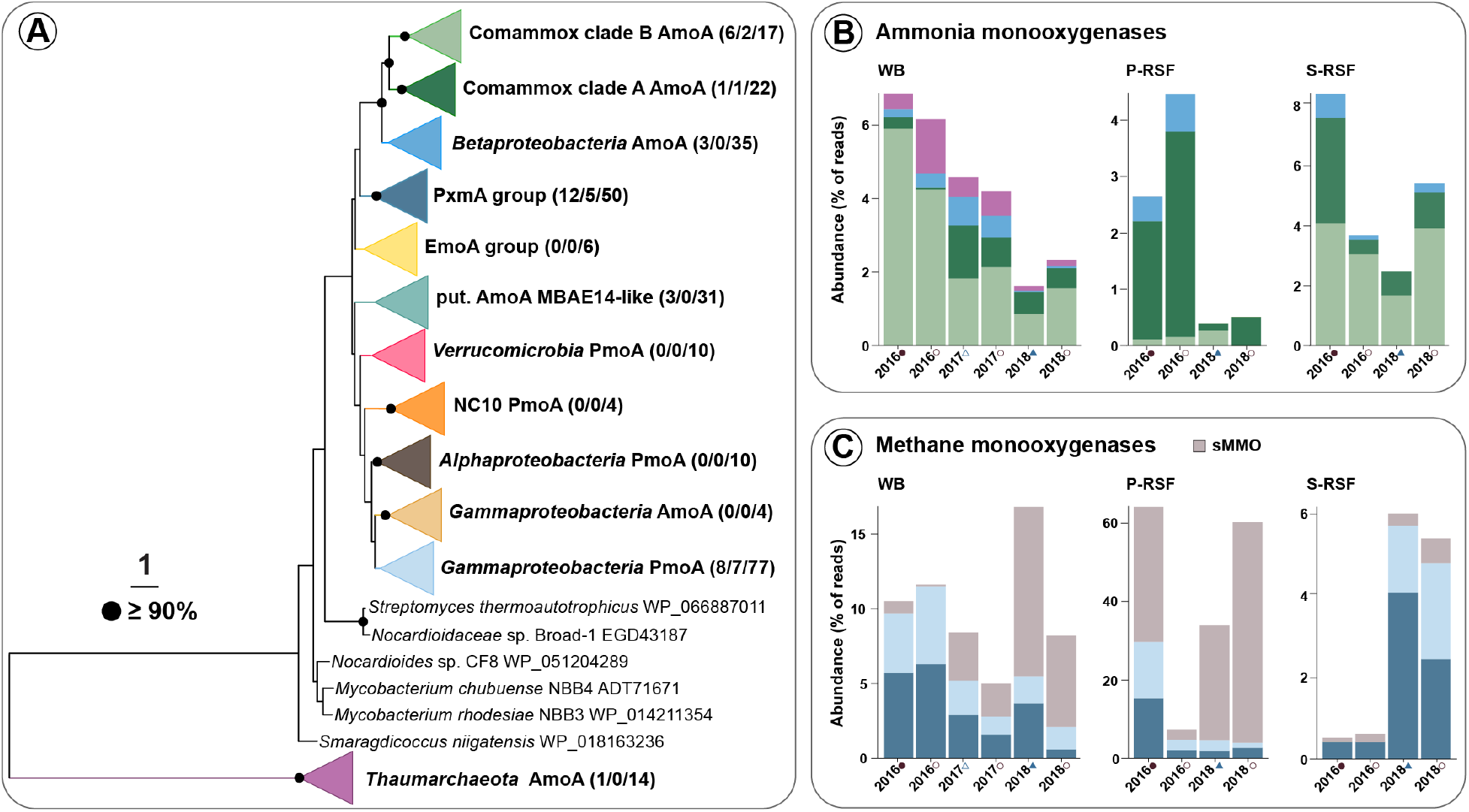
Diversity and abundance of ammonia and methane-oxidizing microorganisms in DWTP Breehei. (**A**) Phylogenetic tree of CuMMO subunit A proteins recovered from the DWTP metagenomes by gene-centric and genome-resolved approaches. Numbers in brackets indicate sequences per metagenome, number of recovered MAGs, and references datasets for each clade, respectively. Amo, ammonia monooxygenase; Emo, ethane monooxygenase; Pmo/Pxm, particulate methane monooxygenase. Bootstrap support values ≥90% are indicated by black dots; the scale bar indicates estimated amino acid substitutions. Normalized abundances of (**B**) ammonia and (**C**) methane oxidizers are shown as proportion of the recovered ammonia and methane monooxygenases, respectively, to the normalized *rpoB* abundance. The *mmoX* gene was used to calculate the abundance of sMMO-containing methanotrophs; most *mmoX* reads were recruited by MAG P-RSF-NP-02 (70-100%). As most available high-quality betaproteobacterial AOB genomes (n=17, comp. ≥90%, number of contigs <5) contain between 2 and 3 *amoA* copies, and MAG P-RSF-NP-02 contained 4 *mmoX* copies, abundances of both genes were normalized for gene copy numbers. Sample labels correspond to Figure 2.

Of the five *Nitrospira* MAGs recovered in this study, one (P-RSF-IL-20) contained *amoA* genes affiliated with clade A, and two (WB-IL-04, WB-IL-16) with clade B (Figure 3A, Table S3). In protein-based phylogenetic analyses, most of the clade A and B comammox AmoA sequences most closely clustered with sequences derived from DWTP metagenomes (Figure S4). Clade A comammox *amoA* genes were detected in all samples from the DWTP Breehei and were the most abundant *amoA* type in P-RSF (0.1-3.6%). In general, ammonia oxidizers were more abundant in 2016 samples than in 2018 (Figure 4B). The clade A comammox *Nitrospira* MAG (P-RSF-IL-20) had the highest coverage in the 2016 P-RSF sample extracted with the Blood & Tissue Kit, while this coverage decreased drastically with another DNA extraction method, indicating a strong extraction bias as discussed above (Table S2). Consequently, the average coverage of this MAG was higher in S-RSF than in P-RSF. In contrast, clade B comammox *amoA* recruited the largest number of reads in WB (0.8-5.9%) and S-RSF samples (1.7-4.1%; Figure 6A), but none of the two clade B-affiliated MAGs was detected in S-RSF, suggesting that not all clade B comammox *Nitrospira* genomes were recovered. The low abundance of clade B comammox *amoA* in P-RSF (Figure 4B) indicates an adaptation of clade B comammox *Nitrospira* to specific niches. In addition to complete nitrifiers, the genus *Nitrospira* also contains canonical nitrite-oxidizing bacteria (NOB) (Daims et al., 2016). Both canonical as well as comammox *Nitrospira* employ the NXR complex for the oxidation of nitrite to nitrate (Figure 3B). Except for one MAG (P-RSF-IL-22), all *Nitrospira* MAGs contained a *nxrAB* gene cluster encoding for the alpha and beta subunit of the NXR complex (Figure 3A, Table S3). MAG P-RSF-IL-22 was of medium quality (74.4% estimated completeness; Table 2) and did not contain any of the genes required for nitrification (Table S3). Consistent with previous studies (Palomo et al., 2019; Poghosyan et al., 2019), phylogenomic analysis using a concatenated alignment of 91 single-copy core genes showed that clade A and B comammox *Nitrospira* formed monophyletic clades within *Nitrospira* lineage II, and all *amoA*-containing MAGs were correctly affiliated with their respective comammox clade (Figure 5A).

**Figure 5.**
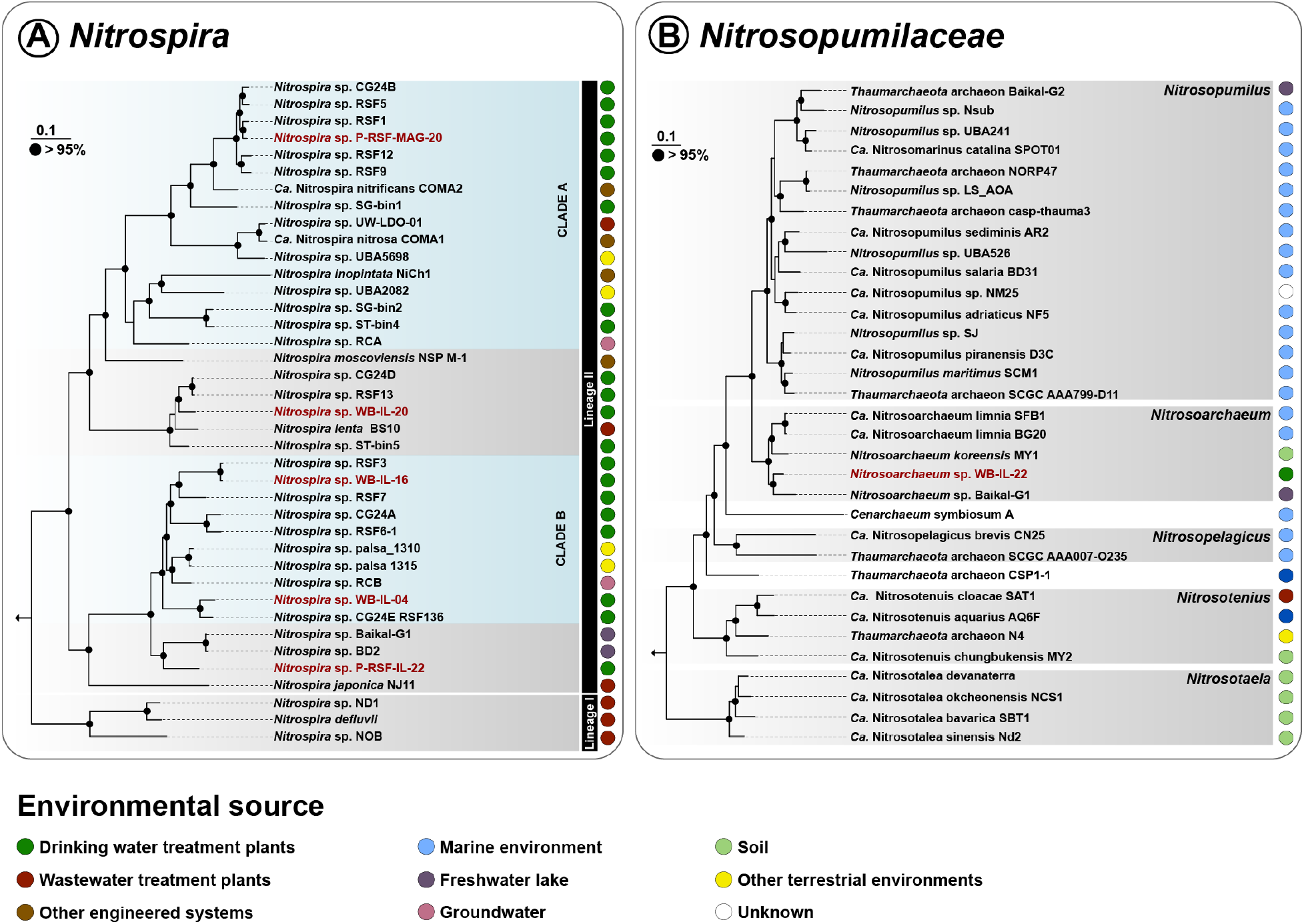
Maximum-likelihood phylogenomic analysis of **(A)** *Nitrospiraceae* based on 91 bacterial single-copy core genes and **(B)** *Nitrosopumilaceae* based on 162 archaeal proteins. DWTP genomes obtained in this study are shown in red. Bootstrap support values ≥95% are indicated by black dots; the scale bars indicate estimated nucleotide **(A)** and amino acid **(B)** substitutions. The positions of the outgroups are indicated by arrows. Environmental origins of the respective genomes are represented by colored circles. Genome names correspond to NCBI RefSeq entries, while in **(B)** classifications at higher taxonomic levels (gray shadings) are according to the GTDB database (Table S4). In **(A)** *Nitrospira* sublineages are delineated by black bars, the comammox clades are highlighted by turquoise shadings.

**Figure 6.**
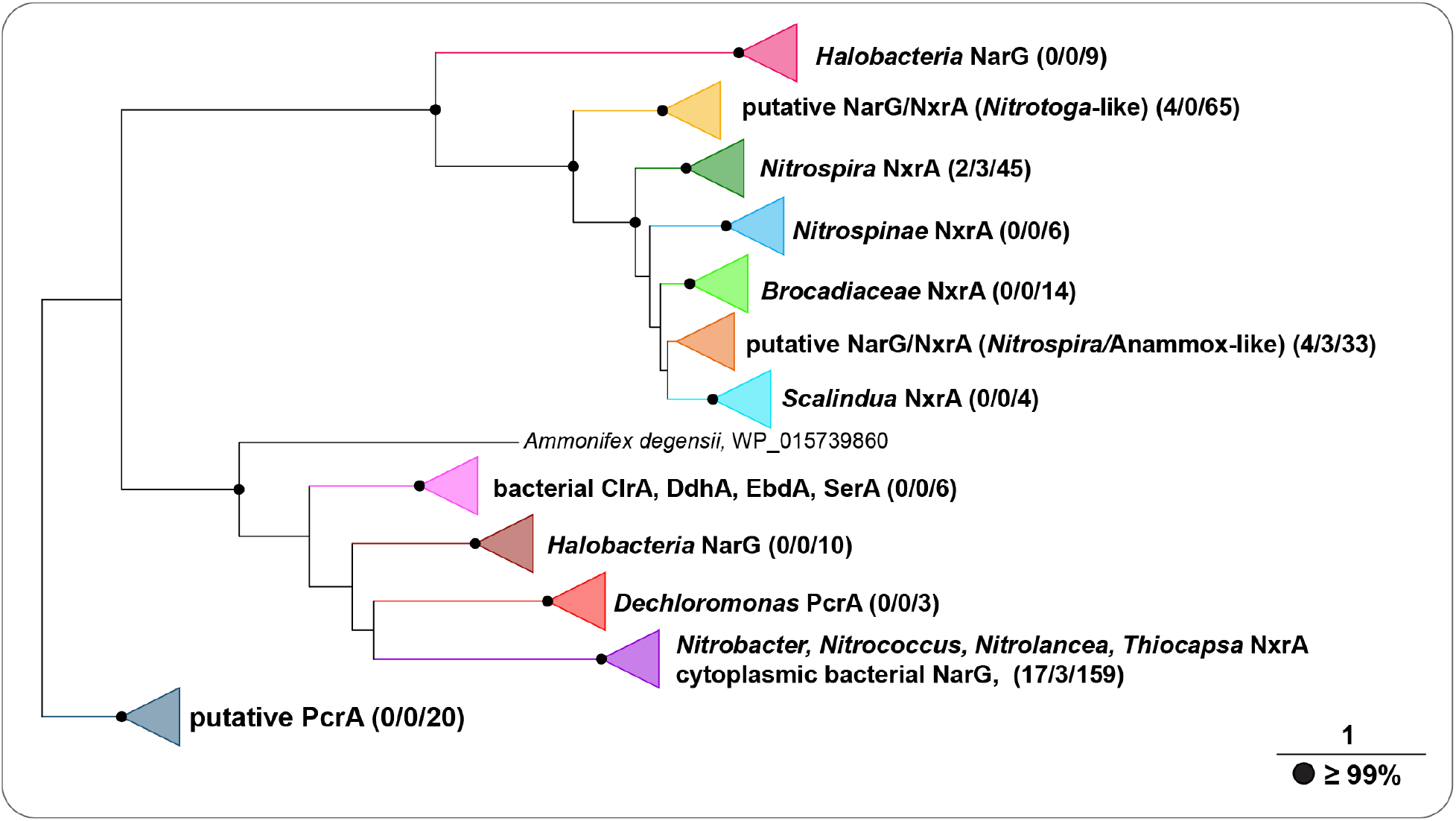
Phylogenetic analysis of NxrA/NarG sequences and related proteins. Numbers in brackets indicate sequences per from DWTP Breehei metagenome, recovered MAGs and reference sequences used for each group. Clr, chlorate reductase; Ddh, dimethylsulfide dehydrogenase; Ebd, ethylbenzene dehydrogenase; Nar, nitrate reductases; Nxr, nitrite oxidoreductase; Pcr, perchlorate reductase; Ser, selenate reductase. Bootstrap support values ≥99% are indicated by black dots; the scale bar indicates estimated amino acid substitutions.

One MAG (WB-IL-22) affiliated with the genus *Nitrosoarchaeum* (AOA; Figure 5B) was retrieved from the WB metagenome; however, it lacked an *amoA* sequence. Since a contig (<1200 bp) containing a *Nitrosoarchaeum*-like *amoA* was identified in the metagenome, the gene most likely is missing from MAG WB-IL-22 due to the size cutoff used during binning (1500 bp). Consistent with gene-based analyses (0.1-1.7%; Figure 4B) the *Nitrosoarchaeum* MAG was found solely in WB, where it accounted for merely ~0.2% of all the assembled reads in WB metagenomes (Figure 2, Table S2). The recovered archaeal AmoA sequences had high similarities to *Nitrosarchaeum koreense* and sequences derived from metagenomic analyses of diverse habitats (Figure S4). Betaproteobacterial *amoA* sequences of members of the genera *Nitrosomonas* and *Nitrosospira* were identified in all samples, but were of low abundance. In addition to the characterized ammonia-oxidizing clades, recently the unclassified Gammaproteobacteria “MBAE14” genome was found to contain putative AMO genes (Mori et al., 2019). From the DWTP Breehei metagenomes, three putative AmoA sequences clustered in this novel sequence group despite low similarities to the putative enzyme of MBAE14 (Figure 4A, S4).

The NXR belongs to the type II dimethyl sulfoxide (DMSO) reductase enzyme family (Lücker et al., 2010; Meincke et al., 1992), which also contains respiratory nitrate reductases (NARs) (Simon and Klotz, 2013). Consequently, many HMM profiling approaches fail to differentiate the two homologous groups. Therefore, a phylogenetic tree was constructed to classify the catalytic alpha subunits (NxrA/NarG) identified in the DWTP metagenomes (Figure 6).

Besides the *Nitrospira* genomes, six additional MAGs contained NXR/NAR gene clusters (Figure 3A). Phylogenetic analysis revealed that the sequences derived from alphaproteobacterial MAGs (P-RSF-IL-15, P-RSF-IL-21, S-RSF-IL-04) were closely affiliated with NarG sequences of known nitrate reducers (Figure S5). The sequences from a MAG affiliated with the *Pedosphaeraceae* (WB-IL-17) and two *Planctomycetota* MAGs (WB-IL-13, S-RSF-IL-06) shared low similarities with known NXRs, but clustered with *Nitrospira, Nitrospina* and anammox bacteria (Figure S5). While the catalytic NxrA subunit of WB-IL-13 contains one Fe-S domain typical for DMSO reductase family molybdoproteins (Lücker et al., 2010), this binding motif for Fe-S clusters is absent from the WB-IL-17 and S-RSF-IL-04 NXR-like proteins. Similar to other periplasmic NXRs (Lücker et al., 2013; Lücker et al., 2010; Spieck et al., 1998), these putative NXRs contain N-terminal twin-arginine motifs for protein translocation into the periplasmic space. Putative genes encoding the NXR gamma subunit (*nxrC*) were found to be co-localized with *nxrAB* in the WB-IL-13 and WB-IL-17 MAGs. Similar to *Nitrospina gracilis* (Lücker et al., 2013) and “*Ca.* Nitrotoga fabula” (Kitzinger et al., 2018), these putative NxrCs contain N-terminal signal peptides necessary for translocation via the Sec pathway. The *Pedosphaeraceae* (WB-IL-17) and one of the *Planctomycetota* MAGs that affiliated with the *Phycisphaerales* (WB-IL-13) were present in WB samples, but of low abundance (Figure 2, Table S1). The second, unclassified *Planctomycetota* MAG (S-RSF-IL-06) was detected only in S-RSF and accounted for 0.5-2% of all assembled reads in the S-RSF metagenomes (Table S1). Additionally, some of the metagenome-derived NXR/NAR sequences were placed into the clade containing the novel NXR type of “*Ca.* Nitrotoga fabula” (Figure S4B) (Kitzinger et al., 2018).

### Methane and one-carbon metabolism

Taxonomic profiling of the extracted 16S rRNA genes (Figure S3) identified *Gammaproteobacteria* as the most dominant taxa in WB (40-55%) and P-RSF (61-67%) samples. On the family level, *Methylomonadaceae* and *Methylophilaceae* formed the most abundant groups in WB and P-RSF (Figure S3B). In total, 14 MAGs belonging to these two phylogenetic groups were recovered, which were mainly present in WB and P-RSF samples (Figure 2).

Phylogenomic analysis of the 14 *Methylomonadaceae* and *Methylophilaceae*-affiliated MAGs using a concatenated alignment of 92 single-copy core genes showed that three of the seven *Methylomonadaceae* MAGs were affiliated with the genus *Methyloglobulus* (Figure 7A). Of the remaining MAGs, two (P-RSF-IL-13 and WB-IL-06) were distantly related to *Methylobacter* and one (WB-IL-19) to *Crenothrix*, while P-RSF-IL-24 clustered separately from the known genera within this family (Figure 7A). Six out of seven *Methylophilaceae* MAGs were affiliated with the genus *Methylotenera,* whereas P-RSF-NP-02 formed a separate branch within the *Methylophilaceae* family (Figure 7B).

**Figure 7.**
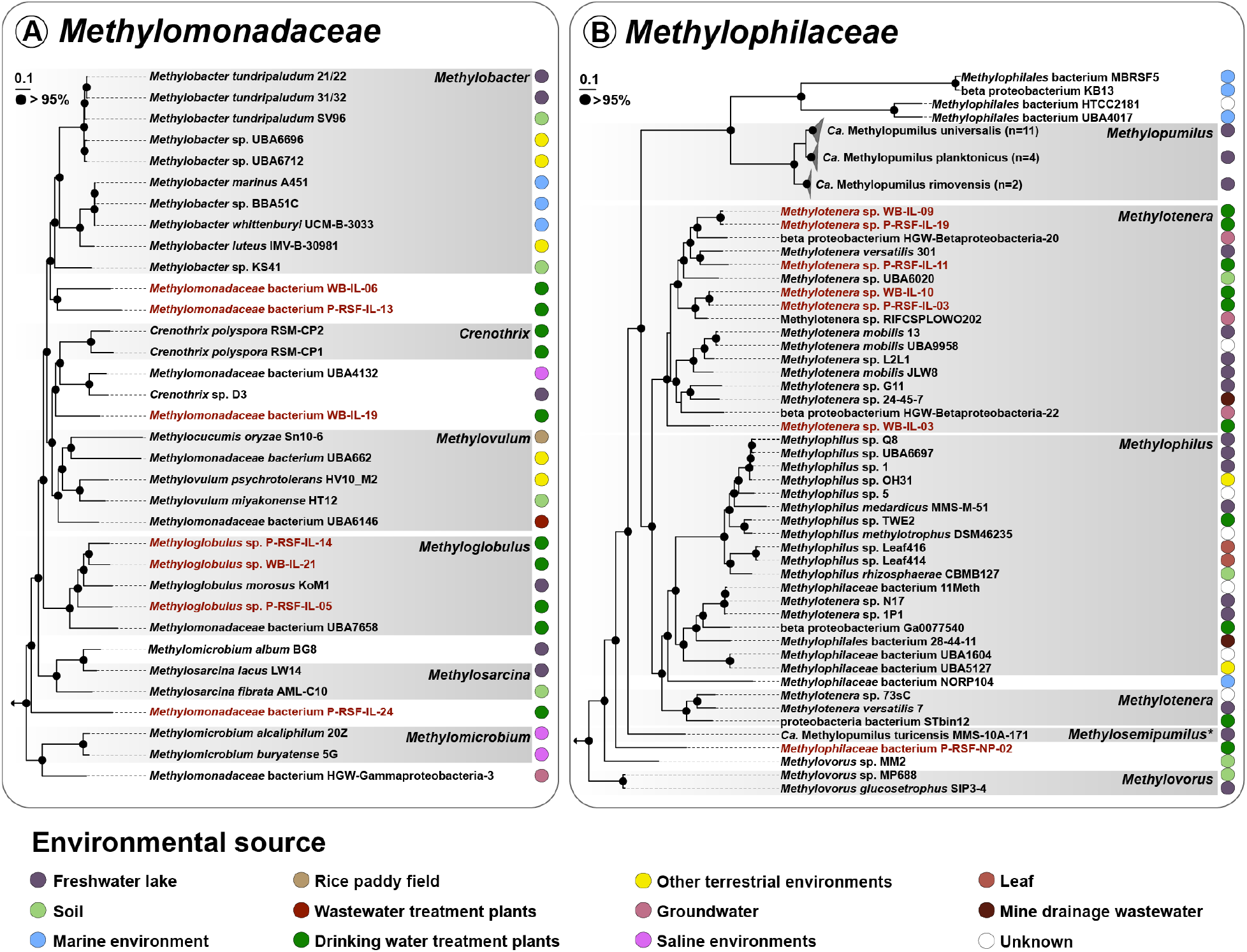
Maximum-likelihood phylogenomic analysis of **(A)** *Methylomonadaceae* and **(B)** *Methylophilaceae* based on 92 bacterial single-copy core genes. MAGs from this study are shown in red. Bootstrap support values ≥95% indicated by black dots; the scale bars indicate estimated nucleotide substitutions. The positions of the outgroups are indicated by arrows. Environmental origins of the respective genomes are represented by colored circles. Genome names correspond to NCBI RefSeq entries, while classifications at higher taxonomic levels are based on GTDB classification (Table S4).

In P-RSF and WB metagenomes, the relative coverage of the *Methylophilaceae* MAGs was much higher than for the *Methylomonadaceae* MAGs (Figure 2, Table S1). These results are consistent with 16S rRNA gene profiling, where *Methylophilaceae* constituted the most abundant family in WB (6-27%) and P-RSF (28-47%) samples and disappeared in S-RSF samples (Figure S3). Particularly, the MAG P-RSF-NP-02 had extremely high coverage in WB (1.6-179×) and P-RSF (27-744×) metagenomes (Table S1). Based on 16S rRNA gene similarity, “*Ca.* Methylosemipumilus turicensis” MMS-10A-171 (Salcher et al., 2019) is the closest described relative of P-RSF-NP-02 (95.7% sequence identity). This value is below the similarity cutoff for species delineation (Stackebrandt and Ebers, 2006), which, in combination with the distinct clustering in the phylogenomic analysis (Figure 7B), indicates that P-RSF-NP-02 probably belongs to a novel genus within the family *Methylophilaceae*. The estimated genomic average nucleotide identity value (ANI; 77.26%) also indicated a novel species distinct from “*Ca.* M. turicensis”. The Breehei DWTP metagenomes and the recovered MAGs furthermore were screened for genes encoding the soluble (sMMO) or particulate (pMMO) methane monooxygenases (Table S3). Notably, a complete operon encoding sMMO *(mmoX1X2YBZDC)* was identified in the highly-covered *Methylophilaceae* MAG P-RSF-NP-02 (Figure 3, Table S3). This was surprising as no previously described member of the family *Methylophilaceae* contains any methane monooxygenase (Salcher et al., 2019). P-RSF-NP-02 also harbors two additional copies of the *mmoX* gene encoding the alpha subunit of the sMMO (Table S3). The two orphan *mmoX* copies share high sequence identities on amino acid level with the operonal *mmoX1* (99.8-100%), while *mmoX2* has ~97% identity to *mmoX1* and the two orphan *mmoX*. A BLAST search against the NCBI RefSeq database identified that the MmoX copies of P-RSF-NP-02 share 83-84% similarity with the *Methylomicrobium buryatense* proteins (Kaluzhnaya et al., 2001). The abundance of methanotrophs in the total community was estimated based on *pmoA, pxmA* and *mmoX* gene coverages (Figure 4C). The sMMO-containing methanotrophs were present at high abundances in the 2018 WB (6-11%) and P-RSF (2.5-56%) samples (Figure 4C), where MAG P-RSF-NP-02 also dominated the total microbial communities (Figure 2). Other samples were dominated by pMMO containing bacteria (Figure 4C.) The gene cluster encoding the structural pMMO subunits *(pmoCAB)* was identified in all *Methylomonadaceae* genomes. Furthermore, five MAGs also harbor the highly divergent *pxmABC* gene cluster encoding for pXMO (Figure 3) (Tavormina et al., 2011). Phylogenetic analysis of the pMMO alpha subunits (PmoA/PxmA) revealed that all sequences recovered in this study, including the metagenome-derived ones, were affiliated with gammaproteobacterial methanotrophs (Figure S4). Besides pMMO, the three *Methylomonadaceae* MAGs (WB-IL-19, P-RSF-IL-13, P-RSF-IL-24) also possess sMMO (Figure 3, Table S3).

In both methanotrophs and non-methanotrophic methylotrophs, methanol oxidation to formaldehyde is catalyzed by pyrroloquinoline quinone-dependent (PQQ) methanol dehydrogenases (MDH) (Keltjens et al., 2014). In this study, the rare earth element-dependent MDH (Ln-MDH) was detected in all *Methylomonadaceae and Methylophilaceae*, in one *Deltaproteobacteria* (S-RSF-IL-11), and one *Hyphomicrobium* (P-RSF-NP-07) MAGs. In contrast, the calcium-dependent MDH (Ca-MDH) was found only in some MAGs (Figure 3A, Table S2). Thus, all analyzed methanotrophs and methylotrophs can oxidize methanol either to formaldehyde by Ca-MDH, or directly to formate by Ln-MDH (Figure 3B) (Pol et al., 2014). Four formaldehyde-oxidizing pathways were identified, including oxidation by single aldehyde dehydrogenases, as well as via tetrahydromethanopterin (H_4_MPT), tetrahydrofolate (H_4_F), or glutathione (GSH)-linked pathways (Figure 3). All the MAGs contained formate dehydrogenases (FDHs), necessary for the oxidation of formate to carbon dioxide.

## Discussion

### DWTP microbiome

The DWTP Breehei microbiome was followed over a time period of three years and characterized using genome-resolved and gene-centric metagenomic approaches. The microbial community structure was studied for the filter material of two sequential rapid sand filters and the biofilm formed on the walls of the primary sand filter (Figure 1). Similar to some Danish DWTPs (Gülay et al., 2016), the location within the DWTP had a strong influence on the microbial community composition. In total, 56 dereplicated near-complete MAGs were recovered, comprising 23, 64, and 14% of the total assembled reads for WB, P-RSF, and S-RSF respectively (Table S1). These MAGs expand our knowledge on the genomic inventory of the main microorganisms involved in contaminant removal from groundwater to produce drinking water. The assembly statistics also indicate that the obtained metagenomic information especially for WB and S-RSF only covers a part of the diversity. In general, the genome-centric approach is a powerful tool to analyze the functional potential of an environmental sample based on recovered MAGs. However, due to low abundance or strain diversity, it often is difficult to obtain good-quality genomes for many microorganisms (Sczyrba et al., 2017). Thus, to examine full ammonia and methane oxidation potential, we also used a gene-centric approach (see below).

In groundwater, iron exists in the ferrous [Fe(II)] and ferric [Fe(III)] state (Chapelle, 2001) and its oxidation can cause severe operational problems during drinking water production, including bad taste, discoloration, staining, deposition in distribution systems leading to aftergrowth and incidences of high turbidity (Emerson and De Vet, 2015; Sharma et al., 2005). Water quality monitoring at the DWTP Breehei over 16 years indicates 99% iron removal efficiency (Table 1). Under oxic conditions and circumneutral pH, iron oxidation occurs both chemically and biologically in these systems (Tekerlekopoulou et al., 2006). Biological iron oxidation is mediated by chemolithoautotrophic microorganisms that obtain energy from oxidizing ferrous iron (Emerson and De Vet, 2015). However, the absence of an universal metabolic pathway for iron oxidation makes it challenging to identify iron-oxidizing bacteria (Emerson and De Vet, 2015). Here, we obtained four MAGs classified as *Gallionella* and *Sideroxydans* species (Figure 2), which are generally regarded as iron-oxidizing bacteria (Bruun et al., 2010; de Vet et al., 2011; Druschel et al., 2008). These microorganisms were found in WB and P-RSF (Figure 2), where the iron load was high (Table 1). It thus can be assumed that *Gallionellaceae* members are the main drivers of biological iron oxidation in this DWTP.

The presence of reduced sulfur compounds such as H_2_S, methanethiol, and dimethyl sulfide in anoxic groundwater adds yet another level of complexity to the system (Emerson and De Vet, 2015; Sercu et al., 2005). These compounds serve as electron donors for sulfur-oxidizing microorganisms. For sulfide oxidation, sulfide:quinone oxidoreductases (SQR) as well as flavocytochrome *c* sulfide dehydrogenases (FCC) were detected in many of the recovered MAGs (Table S3). However, as sulfide reacts with cytochromes, hemeproteins, and other iron-containing compounds, these enzymes are used for detoxification by many bacteria (Cherney et al., 2010; Marcia et al., 2009; Shahak and Hauska, 2008; Zhang et al., 2013). The MAGs WB-IL-01 and P-RSF-IL-21, classified as *Sulfuriferulla* and *Hyphomicrobium*, respectively, contained the heterodisulfide reductase (HdrCBAHypHdrCB) and sulfur oxidation (SoxXYZAB) enzyme complexes, as well as sulfite oxidase (SoeABC; Table S3). The Sox-Hdr-Soe system is a novel pathway found in some chemolithoautotrophic sulfur-oxidizing bacteria and is involved in volatile organosulfur and inorganic sulfur compound degradation (Koch and Dahl, 2018; Watanabe et al., 2019), and also earlier studies provided evidence that the Hdr complex is involved in sulfur oxidation (Boughanemi et al., 2016; Quatrini et al., 2009). Furthermore, recently it has also been shown in *Hyphomicrobium* that the Hdr complex is essential for oxidation of thiosulfate to sulfate (Koch and Dahl, 2018). The Sox-Hdr-Soe system is found in many *Sulfuriferulla* genomes and SoeABC are suggested to be crucial in sulfur oxidizers that lack SoxCD (Watanabe et al., 2019).

Some microbial groups, including *Chloroflexota, Acidobacteriota, Bacteroidota,* and *Myxococcota,* which are originating from groundwater habitats (Griebler and Lueders, 2009; Wegner et al., 2019) are also frequently identified in DWTPs (Albers et al., 2015; Gülay et al., 2016; Palomo et al., 2016). However, we have limited knowledge about their potential role in drinking water treatment. Four MAGs affiliated with *Acidobacteriota* were dominating the microbial community in S-RSF (Figure 2, Figure S3). *Acidobacteriota* are often found in soil under substrate-limited conditions (Hartmann et al., 2015; Jones et al., 2009; Navarrete et al., 2015; Ward et al., 2009). As discussed by Palomo and colleagues (Palomo et al., 2016), *Acidobacteriota* may be involved in carbon cycling in DWTPs. Members of the candidate phyla radiation (CPR) (Brown et al., 2015; Hug et al., 2016) were found only in WB samples (Figure S3). CPR members constitute a substantial fraction of the bacterial diversity (Castelle et al., 2018) and are widely distributed in many environments, including groundwater (Brown et al., 2015; Herrmann et al., 2019; Wegner et al., 2019; Yan et al., 2020) and DWTPs (Bautista-de los Santos et al., 2016; Bruno et al., 2017). One MAG affiliated with “*Ca.* Gracilibacteria” (WB-IL_23) was highly abundant in the WB2018 sample (1-4% of assembled reads) and might have the ability to adapt and survive under low nutrient conditions (Wegner et al., 2019). In groundwater, *Acidobacteriota* and CPR bacteria often co-occur with autotrophic microorganisms (Herrmann et al., 2019). Similar to interactions of nitrifying and heterotrophic bacteria in activated sludge (Okabe et al., 2005), the organic compounds produced by autotrophic bacteria may serve as substrates for *Acidobacteriota* and “*Ca.* Gracilibacteria” in DWTPs.

### Ammonia and nitrite metabolism

The removal of ammonium and nitrite is a vital step in drinking water treatment. Although the drinking water produced in DWTP Breehei is of high quality and free of any nitrogen compounds, intermediate nitrite accumulation is observed in the effluent of the P-RSF, which can be caused by incomplete nitrification (de Vet et al., 2012; Wagner et al., 2016). Thus, profound insights into the microbial key players and their metabolic capabilities and limitations are crucial for optimizing and stabilizing N removal in these systems. The genomic potential for nitrification was mainly identified in MAGs affiliated with the genus *Nitrospira,* which however did not display as high abundances as observed in other DWTP systems (Albers et al., 2015; Gülay et al., 2016; Palomo et al., 2016). Comammox *Nitrospira* were the most abundant nitrifying guild in all DWTP Breehei samples (Figure 4B), corroborating previous findings of *Nitrospira* dominating the nitrifying microbial community in DWTPs (Albers et al., 2015; Gülay et al., 2016). Until recently, the dominance of this group was puzzling since *Nitrospira* were always regarded as strict nitrite-oxidizing bacteria. By now, this imbalance in abundance of *Nitrospira* and canonical AOB can be explained by the presence of complete nitrifying *Nitrospira* in many sand filtration systems (Palomo et al., 2018; Pinto et al., 2015; Wang et al., 2017). Notably, complete nitrifiers are the most abundant nitrifying group in several Danish RSFs (Fowler et al., 2018) and were identified as key drivers of ammonia and nitrite oxidation in these systems (Gülay et al., 2019). Consistent with these previous results, comammox *Nitrospira* also apparently outcompeted canonical AOB and AOA in DWTP Breehei. Comammox clade A was found to dominate in P-RSF samples and clade B in WB and S-RSF (Figure 4B). Thus, sampling location substantially influenced the abundance of the two comammox clades, indicating potential niche partitioning between them. The habitat preferences by different nitrifiers might be explained by their ammonia oxidation kinetics and substrate affinities (Kits et al., 2017; Martens-Habbena et al., 2009; Prosser and Nicol, 2012). Based on the kinetic theory of optimal pathway length (Costa et al., 2006) and physiological analyses of *N. inopinata* (Kits et al., 2017), comammox *Nitrospira* are K-strategists that have a competitive advantage in environments with very low ammonium fluxes. However, since no comammox clade B enrichment culture is currently available, we can only speculate about the niche defining metabolic capabilities of this group. A recent study reported that nitrification activity in forest and paddy soils when subjected to ammonium limitation is associated with clade B rather than clade A comammox *Nitrospira* (Wang et al., 2019). Comammox clade B appeared to dominate also in forest soil under increasing nitrogen load and decreasing pH (Shi et al., 2018). Under acidic conditions, ammonia (NH_3_) is increasingly protonated to ammonium (NH_4_^+^), resulting in extremely low concentrations of bioavailable ammonia, the substrate of AMO. Correspondingly, also in Danish RSFs with influent ammonium concentrations ranging from 0.01-0.53 mg-N/L, clade B constituted up to 75% of the total comammox *Nitrospira* population (Fowler et al., 2018). The higher abundance of clade B comammox *Nitrospira* in S-RSF compared to P-RSF observed here also suggests their adaptation to ammonium-depleted environments. These results indicate that clade B comammox *Nitrospira* may exhibit an even lower half-saturation constant (K_s_) and higher substrate affinities than clade A species and thrive at extremely low ammonium concentrations. However, it will be required to obtain clade B comammox *Nitrospira* in culture to ascertain the physiology of these enigmatic bacteria. In addition to comammox *Nitrospira*, also two canonical *Nitrospira* MAGs were retrieved from the Breehei DWTP metagenomes, indicating that these nitrite oxidizers can interact with canonical ammonia oxidizers as well as with complete nitrifiers. Moreover, we identified MAGs affiliated with the *Verrucomicrobiota* and *Planctomycetota* phyla (Figure 3) that contained NXR-like sequences with similarity to the enzymes of known NOBs. In phylogenetic analyses of NxrA, they clustered with *Nitrospira*, *Nitrospina,* and anammox bacteria (Figure S4B). The apparent periplasmic orientation of the NxrA and the lack of transmembrane helices in the NxrC subunit suggests that the NXR of these putative NOBs may be soluble, as has also been proposed for *Nitrospina gracilis* (Lücker et al., 2013) and “*Ca.* Nitrotoga fabula” (Kitzinger et al., 2018). However, further studies are needed to analyze the potential nitrite-oxidizing capacity of these bacteria. Although in a very low abundances, *Nitrobacter* species are also detected in some RSFs (Tatari et al., 2017), which we however did not observe in DWTP Breehei.

### Methane and one-carbon metabolism

In DWTPs, methane stripping via aeration is preferred over bacterial methane oxidation since microbial activity and growth cause accumulation of extracellular polymeric substances that lead to clogging of the biofilter material (Streese and Stegmann, 2003). However, both WB and P-RSF samples showed high methane-oxidizing capacity (Figure S1). Especially the wall biofilm might counteract methane blowout by oxidizing methane before it leaves the filter and thus reduce methane emissions to the atmosphere. In the global carbon cycle, methane-oxidizing microorganisms play a significant role (Cicerone and Oremland, 1988) as they represent the only known biological methane sink (Aronson et al., 2013). These organisms are able to grow with methane as sole carbon and energy source. The first step of methane oxidation, methane activation and conversion to methanol, is catalyzed either by soluble (sMMO) or particulate (pMMO) methane monooxygenases (Tavormina et al., 2011; Trotsenko and Murrell, 2008). Especially the sMMP is not universal to methanotrophs and only certain phylogenetic groups are known to encode this methane monooxygenase type (Verbeke et al., 2019), usually together with pMMO.

Several studies have shown that the majority of the methane-oxidizing bacteria colonizing the granular material of RSFs are affiliated with the gammaproteobacterial *Methylococcaceae* family (Albers et al., 2015; Gülay et al., 2016; Palomo et al., 2016). Recently, this family was reclassified and split into the *Methylomonadaceae, Methylococcaceae,* and *Methylothermaceae* (Parks et al., 2017). Especially members of the *Methylomonadaceae* have been found in many natural and engineered systems (Flynn et al., 2016; Hoefman et al., 2014; Kalyuzhnaya et al., 2015; Kits et al., 2013; Ogiso et al., 2012; Oswald et al., 2017; Parks et al., 2017; Svenning et al., 2011). In this study, all recovered *Methylomonadaceae* MAGs contained pMMO, and some of them additionally encoded sMMO (Figure 3, Table S2). In addition, several methanotrophic MAGs also contained the highly divergent pXMO enzyme complex encoded by the *pxmABC* gene cluster (Tavormina et al., 2011). Previous studies have shown that pXMO is involved in methane oxidation under hypoxic denitrifying conditions in *Methylomonadaceae* strains (Kits et al., 2015a; Kits et al., 2015b), but its exact function in the sand filters remains to be determined.

Methane oxidation results in the production of various C1 intermediates including methanol and formate, which can be used as substrates by methylotrophic bacteria. In DWTP Breehei, the P-RSF samples were dominated by members of the *Methylophilaceae* family, which was assumed to accommodate only methylotrophic bacteria incapable of growth on methane. Surprisingly, one of the *Methylophilaceae* MAGs (P-RSF-NP-02) harbored a complete gene operon encoding a sMMO (Figure 3, Table S2), indicating a methanotrophic potential. This finding is substantiated by an earlier study, which demonstrated that a member of methylotrophic genus *Methyloceanibacter* can become methanotrophic by acquiring sMMO (Vekeman et al., 2016). The phylogenomic analysis and low ANI value to other members of this family (77.26%) suggest that P-RSF-NP-02 represents a novel species within the *Methylophilaceae* (Figure 7B) and the extremely high coverage of this MAG indicates a major role in methane removal in the DWTP Breehei. The high abundance of this potential new novel methane-oxidizing bacterium in the P-RSF will be facilitated by the high iron content of the influent water, as the sMMO contains a diiron cluster in the active site (Jasniewski and Que, 2018; Wallar and Lipscomb, 1996).

## Conclusions

- The metagenomic analyses enabled us to identify key microbial populations involved in the removal of ammonium and methane. The location within the DWTP Breehei was the most influential factor shaping the microbial community.
- Clade A comammox *Nitrospira* dominated the nitrifying microbial community in P-RSF, while clade B was most abundant in S-RSF where ammonium concentrations are the lowest and, in the biofilm (WB), which is a predicted niche for comammox bacteria.
- The methanotrophic community was dominated by sMMO-containing bacteria, particularly by one novel *Methylophilaceae* member, which might be facilitated by a high iron concentration in the groundwater.

## Author contributions

LP and HOdC collected and processed samples. LP and HK analyzed and interpreted the data. JF and GC contributed to bioinformatics analyses. TvA and GC performed Illumina and Nanopore sequencing. HOdC, MJ and MAHJvK were involved in project discussion and data interpretation. SL conceived the research project. LP, HK, and SL wrote the manuscript with input from all the authors.

## Data availability

The genome sequences of the 56 MAGs recovered in this study and the raw sequencing data have been deposited in GenBank under BioProject number PRJNA622654.

## Acknowledgments

We would like to thank Weren de Vet, Geert Gielens and Kay Bouts for providing necessary information about DWTP Breehei performance and sampling assistance, and Linnea Kop for fruitful discussions. Financial support was provided by the Netherlands Organization for Scientific Research (NWO Talent Programme grants VI.Veni.192.086, 016.Veni.192.062 and 016.Vidi.189.050, and Gravitation grants 024.002.001 [NESSC] and 024.002.002 [SIAM]) and the European Research Council (ERC Advanced Grants 339880 [Ecomom] and 669371 [VOLCANO]).

## References

Albers, C.N., Ellegaard-Jensen, L., Harder, C.B., Rosendahl, S., Knudsen, B.E., Ekelund, F. and Aamand, J. 2015. Groundwater Chemistry Determines the Prokaryotic Community Structure of Waterworks Sand Filters. Environ Sci Technol 49(2), 839–846.

Alneberg, J., Bjarnason, B.S., de Bruijn, I., Schirmer, M., Quick, J., Ijaz, U.Z., Lahti, L., Loman, N.J., Andersson, A.F. and Quince, C. 2014. Binning metagenomic contigs by coverage and composition. Nat Methods 11(11), 1144–1146.

Armenteros, J.J.A., Tsirigos, K.D., Sonderby, C.K., Petersen, T.N., Winther, O., Brunak, S., von Heijne, G. and Nielsen, H. 2019. SignalP 5.0 improves signal peptide predictions using deep neural networks. Nat Biotechnol 37(4), 420–423.

Aronson, E.L., Allison, S.D. and Helliker, B.R. 2013. Environmental impacts on the diversity of methane-cycling microbes and their resultant function. Front Microbiol (4:225).

Bautista-de los Santos, Q.M., Schroeder, J.L., Sevillano-Rivera, M.C., Sungthong, R., Ijaz, U.Z., Sloan, W.T. and Pinto, A.J. 2016. Emerging investigators series: microbial communities in full-scale drinking water distribution systems - a meta-analysis. Environ Sci-Wat Res Technol 2(4), 631–644.

Beech, W.B. and Sunner, J. 2004. Biocorrosion: towards understanding interactions between biofilms and metals. Curr Opin Biotechnol 15(3), 181–186.

Boughanemi, S., Lyonnet, J., Infossi, P., Bauzan, M., Kosta, A., Lignon, S., Giudici-Orticoni, M.T. and Guiral, M. 2016. Microbial oxidative sulfur metabolism: biochemical evidence of the membrane-bound heterodisulfide reductase-like complex of the bacterium Aquifex aeolicus. FEMS Microbiol Lett 363(15).

Bowers, R.M., Kyrpides, N.C., Stepanauskas, R., Harmon-Smith, M., Doud, D., Reddy, T.B.K., Schulz, F., Jarett, J., Rivers, A.R., Eloe-Fadrosh, E.A., Tringe, S.G., Ivanova, N.N., Copeland, A., Clum, A., Becraft, E.D., Malmstrom, R.R., Birren, B., Podar, M., Bork, P., Weinstock, G.M., Garrity, G.M., Dodsworth, J.A., Yooseph, S., Sutton, G., Glockner, F.O., Gilbert, J.A., Nelson, W.C., Hallam, S.J., Jungbluth, S.P., Ettema, T.J.G., Tighe, S., Konstantinidis, K.T., Liu, W.T., Baker, B.J., Rattei, T., Eisen, J.A., Hedlund, B., McMahon, K.D., Fierer, N., Knight, R., Finn, R., Cochrane, G., Karsch-Mizrachi, I., Tyson, G.W., Rinke, C., Lapidus, A., Meyer, F., Yilmaz, P., Parks, D.H., Eren, A.M., Schriml, L., Banfield, J.F., Hugenholtz, P., Woyke, T. and Genome Stand, C. 2017. Minimum information about a single amplified genome (MISAG) and a metagenome-assembled genome (MIMAG) of bacteria and archaea. Nat Biotechnol 35(8), 725–731.

Brown, C.T., Hug, L.A., Thomas, B.C., Sharon, I., Castelle, C.J., Singh, A., Wilkins, M.J., Wrighton, K.C., Williams, K.H. and Banfield, J.F. 2015. Unusual biology across a group comprising more than 15% of domain Bacteria. Nature 523(7559), 208–211.

Bruno, A., Sandionigi, A., Rizzi, E., Bernasconi, M., Vicario, S., Galimberti, A., Cocuzza, C., Labra, M. and Casiraghi, M. 2017. Exploring the under-investigated "microbial dark matter" of drinking water treatment plants. Sci Rep 7, 44350.

Bruun, A.M., Finster, K., Gunnlaugsson, H.P., Nornberg, P. and Friedrich, M.W. 2010. A Comprehensive Investigation on Iron Cycling in a Freshwater Seep Including Microscopy, Cultivation and Molecular Community Analysis. Geomicrobiol J 27(1), 15–34.

Campbell, J.H., O’Donoghue, P., Campbell, A.G., Schwientek, P., Sczyrba, A., Woyke, T., Soll, D. and Podar, M. 2013. UGA is an additional glycine codon in uncultured SR1 bacteria from the human microbiota. Proc Natl Acad Sci USA 110(14), 5540–5545.

Camper, A.K. 2004. Involvement of humic substances in regrowth. Int J Food Microbiol 92(3), 355–364.

Castelle, C.J., Brown, C.T., Anantharaman, K., Probst, A.J., Huang, R.H. and Banfield, J.F. 2018. Biosynthetic capacity, metabolic variety and unusual biology in the CPR and DPANN radiations. Nat Rev Microbiol 16(10), 629–645.

Chapelle, F.H. (2001) Ground-water Microbiology and Geochemistry, John Wiley & Sons, New York.

Cherney, M.M., Zhang, Y.F., Solomonson, M., Weiner, J.H. and James, M.N.G. 2010. Crystal Structure of Sulfide:Quinone Oxidoreductase from Acidithiobacillus ferrooxidans: Insights into Sulfidotrophic Respiration and Detoxification. J Mol Biol 398(2), 292–305.

Cicerone, R.J. and Oremland, R.S. 1988. Biogeochemical aspects of atmospheric methane. Global Biogeochem Cycles 2(4), 299–327.

Costa, E., Pérez, J. and Kreft, J.U. 2006. Why is metabolic labour divided in nitrification? Trends Microbiol. 14(5), 213–219.

de Vet, W., Dinkla, I.J.T., Rietveld, L.C. and van Loosdrecht, M.C.M. 2011. Biological iron oxidation by Gallionella spp. in drinking water production under fully aerated conditions. Water Res 45(17), 5389–5398.

de Vet, W., van Loosdrecht, M.C.M. and Rietveld, L.C. 2012. Phosphorus limitation in nitrifying groundwater filters. Water Res 46(4), 1061–1069.

Delmont, T.O. and Eren, A.M. 2016. Identifying contamination with advanced visualization and analysis practices: metagenomic approaches for eukaryotic genome assemblies. PeerJ 4, e1839.

Druschel, G.K., Emerson, D., Sutka, R., Suchecki, P. and Luther, G.W. 2008. Low-oxygen and chemical kinetic constraints on the geochemical niche of neutrophilic iron(II) oxidizing microorganisms. Geochim Cosmochim Acta 72(14), 3358–3370.

EC 2016 Synthesis Report on the Quality of Drinking Water in the Union examining Member States’ reports for the 2011-2013 period, foreseen under Article 13(5) of Directive 98/83/EC. 2016, https://ec.europa.eu/environment/water/water-drink/reporting_en.html.

Eddy, S.R. 2011. Accelerated Profile HMM Searches. PLoS Comput Biol 7(e1002195).

Emerson, D. and De Vet, W. 2015. The Role of FeOB in Engineered Water Ecosystems: A Review. J Am Water Works Assn 107(1), E47–E57.

Eren, A.M., Esen, O.C., Quince, C., Vineis, J.H., Morrison, H.G., Sogin, M.L. and Delmont, T.O. 2015. Anvi’o: an advanced analysis and visualization platformfor ‘omics data. PeerJ 3, e1319.

Fish, J.A., Chai, B.L., Wang, Q., Sun, Y.N., Brown, C.T., Tiedje, J.M. and Cole, J.R. 2013. FunGene: the functional gene pipeline and repository. Front Microbiol 4, 291.

Flynn, J.D., Hirayama, H., Sakai, Y., Dunfield, P.F., Klotz, M.G., Knief, C., Op den Camp, H.J.M., Jetten, M.S.M., Khmelenina, V.N., Trotsenko, Y.A., Murrell, J.C., Semrau, J.D., Svenning, M.M., Stein, L.Y., Kyrpides, N., Shapiro, N., Woyke, T., Bringel, F., Vuilleumier, S., DiSpirito, A.A. and Kalyuzhnaya, M.G. 2016. Draft Genome Sequences of Gammaproteobacterial Methanotrophs Isolated from Marine Ecosystems. Genome Announc 4(1), 2.

Fowler, S.J., Palomo, A., Dechesne, A., Mines, P.D. and Smets, B.F. 2018. Comammox Nitrospira are abundant ammonia oxidizers in diverse groundwater-fed rapid sand filter communities. Environ Microbiol 20(3), 1002–1015.

Griebler, C. and Lueders, T. 2009. Microbial biodiversity in groundwater ecosystems. Freshw Biol 54(4), 649–677.

Gülay, A., Fowler, J., Tatari, K., Thamdrup, B., Albrechtsen, H.-J., Al-Soud, W.A., Sørensen, S.J. and Smets, B.F. 2019. DNA and RNA-SIP reveal Nitrospira spp. as key drivers of nitrification in groundwater-fed biofilters. mBio, 10: e01870–01819.

Gülay, A.M., S., Albrechtsen, H.J., Abu Al-Soud, W., Sorensen, S.J. and Smets, B.F. 2016. Ecological patterns, diversity and core taxa of microbial communities in groundwater-fed rapid gravity filters. ISME J. 10(9), 2209–2222.

Hartmann, M., Frey, B., Mayer, J., Mader, P. and Widmer, F. 2015. Distinct soil microbial diversity under long-term organic and conventional farming. ISME J 9(5), 1177–1194.

Herrmann, M., Wegner, C.E., Taubert, M., Geesink, P., Lehmann, K., Yan, L.J., Lehmann, R., Totsche, K.U. and Kusel, K. 2019. Predominance of Cand. Patescibacteria in Groundwater Is Caused by Their Preferential Mobilization From Soils and Flourishing Under Oligotrophic Conditions. Front Microbiol 10, 1407.

Hoefman, S., Heylen, K. and De Vos, P. 2014. Methylomonas lenta sp nov., a methanotroph isolated from manure and a denitrification tank. Int J Syst Evol Microbiol 64, 1210–1217.

Hug, L.A., Baker, B.J., Anantharaman, K., Brown, C.T., Probst, A.J., Castelle, C.J., Butterfield, C.N., Hernsdorf, A.W., Amano, Y., Ise, K., Suzuki, Y., Dudek, N., Relman, D.A., Finstad, K.M., Amundson, R., Thomas, B.C. and Banfield, J.F. 2016. A new view of the tree of life. Nat Microbiol 1: 16048(5).

Hyatt, D. 2010. Prodigal: prokaryotic gene recognition and translation initiation site identification. BMC Bioinf 11(1), 119.

Jasniewski, A.J. and Que, L. 2018. Dioxygen Activation by Nonheme Diiron Enzymes: Diverse Dioxygen Adducts, High-Valent Intermediates, and Related Model Complexes. Chem Rev 118(5), 2554–2592.

Jones, R.T., Robeson, M.S., Lauber, C.L., Hamady, M., Knight, R. and Fierer, N. 2009. A comprehensive survey of soil acidobacterial diversity using pyrosequencing and clone library analyses. ISME J 3(4), 442–453.

Kalyuzhnaya, M.G., Lamb, A.E., McTaggart, T.L., Oshkin, I.Y., Shapiro, N., Woyke, T. and Chistoserdova, L. 2015. Draft Genome Sequences of Gammaproteobacterial Methanotrophs Isolated from Lake Washington Sediment. Genome Announc 3(2: e00103–15).

Khadka, R., Clothier, L., Wang, L., Lim, C.K., Klotz, M.G. and Dunfield, P.F. 2018. Evolutionary History of Copper Membrane Monooxygenases. Front Microbiol 9, 2493.

Kim, D., Song, L., Breitwieser, F.P. and Salzberg, S.L. 2016. Centrifuge: rapid and sensitive classification of metagenomic sequences. Genome Res 26(12), 1721–1729.

Kits, K.D., Campbell, D.J., Rosana, A.R. and Stein, L.Y. 2015a. Diverse electron sources support denitrification under hypoxia in the obligate methanotroph Methylomicrobium album strain BG8. Front Microbiol 6, 1072.

Kits, K.D., Kalyuzhnaya, M.G., Klotz, M.G., Jetten, M.S., Op den Camp, H.J., Vuilleumier, S., Bringel, F., Dispirito, A.A., Murrell, J.C., Bruce, D., Cheng, J.F., Copeland, A., Goodwin, L., Hauser, L., Lajus, A., Land, M.L., Lapidus, A., Lucas, S., Medigue, C., Pitluck, S., Woyke, T., Zeytun, A. and Stein, L.Y. 2013. Genome Sequence of the Obligate Gammaproteobacterial Methanotroph Methylomicrobium album Strain BG8. Genome Announc 1(2), e0017013.

Kits, K.D., Klotz, M.G. and Stein, L.Y. 2015b. Methane oxidation coupled to nitrate reduction under hypoxia by the Gammaproteobacterium Methylomonas denitrificans, sp nov type strain FJG1. Environmental Microbiology 17(9), 3219–3232.

Kits, K.D., Sedlacek, C.J., Lebedeva, E.V., Han, P., Bulaev, A., Pjevac, P., Daebeler, A., Romano, S., Albertsen, M., Stein, L.Y., Daims, H. and Wagner, M. 2017. Kinetic analysis of a complete nitrifier reveals an oligotrophic lifestyle. Nature 549(7671), 269–272.

Kitzinger, K., Koch, H., Lucker, S., Sedlacek, C.J., Herbold, C., Schwarz, J., Daebeler, A., Mueller, A.J., Lukumbuzya, M., Romano, S., Leisch, N., Karst, S.M., Kirkegaard, R., Albertsen, M., Nielsen, P.H., Wagner, M. and Daims, H. 2018. Characterization of the First "C48" Isolate Reveals Metabolic Versatility and Separate Evolution of Widespread Nitrite-Oxidizing Bacteria. mBio 9(4:e01186–18).

Koch, T. and Dahl, C. 2018. A novel bacterial sulfur oxidation pathway provides a new link between the cycles of organic and inorganic sulfur compounds. ISME J 12(10), 2479–2491.

Koren, S., Walenz, B.P., Berlin, K., Miller, J.R., Bergman, N.H. and Phillippy, A.M. 2017. Canu: scalable and accurate long-read assembly via adaptive k-mer weighting and repeat separation. Genome Res 27(5), 722–736.

Koster, J. and Rahmann, S. 2012. Snakemake-a scalable bioinformatics workflow engine. Bioinformatics 28(19), 2520–2522.

Kowalchuk, G.A., de Bruijn, F.J., Head, I.M., Akkermans, A.D. and van Elsas, J.D. 2004 Molecular microbial ecology manual (MMEM), 2nd ed., vol. 1., Kluwer Academic Publishing, London, United Kingdom.

Krogh, A., Larsson, B., von Heijne, G. and Sonnhammer, E.L.L. 2001. Predicting transmembrane protein topology with a hidden Markov model: Application to complete genomes. J Mol Biol 305(3), 567–580.

Langmead, B. and Salzberg, S.L. 2012. Fast gapped-read alignment with Bowtie 2. Nat Methods 9(4), 357–359.

Li, D.H., Liu, C.M., Luo, R.B., Sadakane, K. and Lam, T.W. 2015. MEGAHIT: an ultra-fast single-node solution for large and complex metagenomics assembly via succinct de Bruijn graph. Bioinformatics 31(10), 1674–1676.

Li, H. 2018. Minimap2: pairwise alignment for nucleotide sequences. Bioinformatics 34(18), 3094–3100.

Li, H. and Durbin, R. 2010. Fast and accurate long-read alignment with Burrows-Wheeler transform. Bioinformatics 26(5), 589–595.

Li, H., Handsaker, B., Wysoker, A., Fennell, T., Ruan, J., Homer, N., Marth, G., Abecasis, G., Durbin, R. and Genome Project Data, P. 2009. The Sequence Alignment/Map format and SAMtools. Bioinformatics 25(16), 2078–2079.

Li, H.S. and Carlson, K.H. 2014. Distribution and Origin of Groundwater Methane in the Wattenberg Oil and Gas Field of Northern Colorado. Environ Sci Technol 48(3), 1484–1491.

Li, Q., Yu, S.L., Li, L., Liu, G.C., Gu, Z.Y., Liu, M.M., Liu, Z.Y., Ye, Y.B., Xia, Q. and Ren, L.M. 2017. Microbial Communities Shaped by Treatment Processes in a Drinking Water Treatment Plant and Their Contribution and Threat to Drinking Water Safety. Front Microbiol 8, 2465.

Ludwig, W., Strunk, O., Westram, R., Richter, L., Meier, H., Yadhukumar, Buchner, A., Lai, T., Steppi, S., Jobb, G., Forster, W., Brettske, I., Gerber, S., Ginhart, A.W., Gross, O., Grumann, S., Hermann, S., Jost, R., Konig, A., Liss, T., Lussmann, R., May, M., Nonhoff, B., Reichel, B., Strehlow, R., Stamatakis, A., Stuckmann, N., Vilbig, A., Lenke, M., Ludwig, T., Bode, A. and Schleifer, K.H. 2004. ARB: a software environment for sequence data. Nucleic Acids Res 32(4), 1363–1371.

Lücker, S., Nowka, B., Rattei, T., Spieck, E. and Daims, H. 2013. The Genome of Nitrospina gracilis Illuminates the Metabolism and Evolution of the Major Marine Nitrite Oxidizer. Front Microbiol 4, 27.

Maksimavičius, E. and Roslev, P. 2020. Methane emission and methanotrophic activity in groundwater fed drinking water treatment plants. Water Supply, ws2020009.

Marcia, M., Ermler, U., Peng, G.H. and Michel, H. 2009. The structure of Aquifex aeolicus sulfide:quinone oxidoreductase, a basis to understand sulfide detoxification and respiration. Proc. Natl. Acad. Sci. U.S.A. 106(24), 9625–9630.

Martens-Habbena, W., Berube, P.M., Urakawa, H., de la Torre, J.R. and Stahl, D.A. 2009. Ammonia oxidation kinetics determine niche separation of nitrifying Archaea and Bacteria. Nature 461(7266), 976–979.

Matsen, F.A., Kodner, R.B. and Armbrust, E.V. 2010. pplacer: linear time maximum-likelihood and Bayesian phylogenetic placement of sequences onto a fixed reference tree. Bmc Bioinf 11(538).

Miller, M.A., Pfeiffer, W. and Schwartz, T. 2010 Creating the CIPRES Science Gateway for inference of large phylogenetic trees. Gateway Computing Environments Workshop (GCE), pp. 1–8.

Na, S.I., Kim, Y.O., Yoon, S.H., Ha, S.M., Baek, I. and Chun, J. 2018. UBCG: Up-to-date bacterial core gene set and pipeline for phylogenomic tree reconstruction. J Microbiol 56(4), 280–285.

Navarrete, A.A., Venturini, A.M., Meyer, K.M., Klein, A.M., Tiedje, J.M., Bohannan, B.J.M., Nusslein, K., Tsai, S.M. and Rodrigues, J.L.M. 2015. Differential Response of Acidobacteria Subgroups to Forest-to-Pasture Conversion and Their Biogeographic Patterns in the Western Brazilian Amazon. Front Microbiol 6, 1443.

Nurk, S., Meleshko, D., Korobeynikov, A. and Pevzner, P.A. 2017. metaSPAdes: a new versatile metagenomic assembler. Genome Res 27(5), 824–834.

Ogiso, T., Ueno, C., Dianou, D., Huy, T.V., Katayama, A., Kimura, M. and Asakawa, S. 2012. Methylomonas koyamae sp nov., a type I methane-oxidizing bacterium from floodwater of a rice paddy field. Int J Syst Evol Microbiol 62, 1832–1837.

Oh, S., Hammes, F. and Liu, W.T. 2018. Metagenomic characterization of biofilter microbial communities in a full-scale drinking water treatment plant. Water Res 128, 278–285.

Okabe, S., Kindaichi, T. and Ito, T. 2005. Fate of 14C-Labeled Microbial Products Derived from Nitrifying Bacteria in Autotrophic Nitrifying Biofilms. Appl Environ Microbiol 71(7), 3987.

Okoniewska, E., Lach, J., Kacprzak, M. and Neczaj, E. 2007. The removal of manganese, iron and ammonium nitrogen on impregnated activated carbon. Desalination 206(1-3), 251–258.

Olm, M.R., Brown, C.T., Brooks, B. and Banfield, J.F. 2017. dRep: a tool for fast and accurate genomic comparisons that enables improved genome recovery from metagenomes through de-replication. ISME J 11(12), 2864–2868.

Osborn, S.G., Vengosh, A., Warner, N.R. and Jackson, R.B. 2011. Methane contamination of drinking water accompanying gas-well drilling and hydraulic fracturing. Proc. Natl. Acad. Sci. U.S.A. 108(20), 8172–8176.

Oswald, K., Graf, J.S., Littmann, S., Tienken, D., Brand, A., Wehrli, B., Albertsen, M., Daims, H., Wagner, M., Kuypers, M.M.M., Schubert, C.J. and Milucka, J. 2017. Crenothrix are major methane consumers in stratified lakes. ISME J. 11(9), 2124–2140.

Palomo, A., Dechesne, A. and Smets, B.F. 2019. Genomic profiling of Nitrospira species reveals ecological success of comammox Nitrospira. bioRxiv, 612226.

Palomo, A., Jane Fowler, S., Gulay, A., Rasmussen, S., Sicheritz-Ponten, T. and Smets, B.F. 2016. Metagenomic analysis of rapid gravity sand filter microbial communities suggests novel physiology of Nitrospira spp. ISME J 10(11), 2569–2581.

Palomo, A., Pedersen, A.G., Fowler, S.J., Dechesne, A., Sicheritz-Ponten, T. and Smets, B.F. 2018. Comparative genomics sheds light on niche differentiation and the evolutionary history of comammox Nitrospira. ISME J. 12(7), 1779–1793.

Parks, D.H., Chuvochina, M., Waite, D.W., Rinke, C., Skarshewski, A., Chaumeil, P.A. and Hugenholtz, P. 2018. A standardized bacterial taxonomy based on genome phylogeny substantially revises the tree of life. Nature Biotechnology 36(10), 996–1004.

Parks, D.H., Imelfort, M., Skennerton, C.T., Hugenholtz, P. and Tyson, G.W. 2015. CheckM: assessing the quality of microbial genomes recovered from isolates, single cells, and metagenomes. Genome research 25(7), 1043–1055.

Parks, D.H., Rinke, C., Chuvochina, M., Chaumeil, P.A., Woodcroft, B., Evans, P.N., Hugenholtz, P. and Tyson, G.W. 2017. Recovery of nearly 8,000 metagenome-assembled genomes substantially expands the tree of life. Nat Microbiol 3(2), 253–253.

Pinto, A.J., Marcus, D.N., Ijaz, U.Z., Santos, Q., Dick, G.J. and Raskin, L. 2015. Metagenomic Evidence for the Presence of Comammox Nitrospira-Like Bacteria in a Drinking Water System. Msphere 1(1), e00054–00015.

Pinto, A.J., Xi, C.W. and Raskin, L. 2012. Bacterial Community Structure in the Drinking Water Microbiome Is Governed by Filtration Processes. Environ Sci Technol 46(16), 8851–8859.

Poghosyan, L., Koch, H., Lavy, A., Frank, J., van Kessel, M.A.H.J., Jetten, M.S.M., Banfield, J.F. and Lücker, S. 2019. Metagenomic recovery of two distinct comammox Nitrospira from the terrestrial subsurface. Environ Microbiol 21(10), 3627–3637.

Pol, A., Barends, T.R.M., Dietl, A., Khadem, A.F., Eygensteyn, J., Jetten, M.S.M. and Op den Camp, H.J.M. 2014. Rare earth metals are essential for methanotrophic life in volcanic mudpots. Environ Microbiol 16(1), 255–264.

Proctor, C.R. and Hammes, F. 2015. Drinking water microbiology - from measurement to management. Curr Opin Biotechnol 33, 87–94.

Prosser, J.I. and Nicol, G.W. 2012. Archaeal and bacterial ammonia-oxidisers in soil: the quest for niche specialisation and differentiation. Trends Microbiol 20(11), 523–531.

Quatrini, R., Appia-Ayme, C., Denis, Y., Jedlicki, E., Holmes, D.S. and Bonnefoy, V. 2009. Extending the models for iron and sulfur oxidation in the extreme Acidophile Acidithiobacillus ferrooxidans. Bmc Genomics 10(394).

Racine, J.S. 2012. RStudio: A Platform-Independent IDE for R and Sweave. J Appl Econ 27(1), 167–172.

Rinke, C., Schwientek, P., Sczyrba, A., Ivanova, N.N., Anderson, I.J., Cheng, J.F., Darling, A., Malfatti, S., Swan, B.K., Gies, E.A., Dodsworth, J.A., Hedlund, B.P., Tsiamis, G., Sievert, S.M., Liu, W.T., Eisen, J.A., Hallam, S.J., Kyrpides, N.C., Stepanauskas, R., Rubin, E.M., Hugenholtz, P. and Woyke, T. 2013. Insights into the phylogeny and coding potential of microbial dark matter. Nature 499(7459), 431–437.

Rittmann, B.E., Tangy, Y., Meyer, K. and Bellamy, W.D. 2012 Biological processes, chapter 17. In: Water treatment design. Ed Randtke SJ and Horsley MB, Am Water Works Assoc (AWWA), McGraw Hill.

Sczyrba, A., Hofmann, P., Belmann, P., Koslicki, D., Janssen, S., Dröge, J., Gregor, I., Majda, S., Fiedler, J., Dahms, E., Bremges, A., Fritz, A., Garrido-Oter, R., Jørgensen, T.S., Shapiro, N., Blood, P.D., Gurevich, A., Bai, Y., Turaev, D., DeMaere, M.Z., Chikhi, R., Nagarajan, N., Quince, C., Meyer, F., Balvočiūtė, M., Hansen, L.H., Sørensen, S.J., Chia, B.K.H., Denis, B., Froula, J.L., Wang, Z., Egan, R., Don Kang, D., Cook, J.J., Deltel, C., Beckstette, M., Lemaitre, C., Peterlongo, P., Rizk, G., Lavenier, D., Wu, Y.-W., Singer, S.W., Jain, C., Strous, M., Klingenberg, H., Meinicke, P., Barton, M.D., Lingner, T., Lin, H.-H., Liao, Y.-C., Silva, G.G.Z., Cuevas, D.A., Edwards, R.A., Saha, S., Piro, V.C., Renard, B.Y., Pop, M., Klenk, H.-P., Göker, M., Kyrpides, N.C., Woyke, T., Vorholt, J.A., Schulze-Lefert, P., Rubin, E.M., Darling, A.E., Rattei, T. and McHardy, A.C. 2017. Critical Assessment of Metagenome Interpretation—a benchmark of metagenomics software. Nat Methods 14(11), 1063–1071.

Seemann, T. 2014. Prokka: rapid prokaryotic genome annotation. Bioinformatics 30(14), 2068–2069.

Sercu, B., Nunez, D., Van Langenhove, H., Aroca, G. and Verstraete, W. 2005. Operational and microbiological aspects of a bioaugmented two-stage biotrickling filter removing hydrogen sulfide and dimethyl sulfide. Biotechnol Bioeng 90(2), 259–269.

Shahak, Y. and Hauska, G. (2008) Sulfur Metabolism in Phototrophic Organisms. Hell, R., Dahl, C., Knaff, D. and Leustek, T. (eds), pp. 319–335, Springer Netherlands, Dordrecht.

Sharma, S.K., Petrusevski, B. and Schippers, J.C. 2005. Biological iron removal from groundwater: a review. J. Water Supply Res. Technol. AQUA 54(4), 239–247.

Shi, X.Z., Hu, H.W., Wang, J.Q., He, J.Z., Zheng, C.Y., Wan, X.H. and Huang, Z.Q. 2018. Niche separation of comammox Nitrospira and canonical ammonia oxidizers in an acidic subtropical forest soil under long-term nitrogen deposition. Soil Biol Biochem 126, 114–122.

Stamatakis, A. 2014. RAxML version 8: a tool for phylogenetic analysis and post-analysis of large phylogenies. Bioinformatics 30(9), 1312–1313.

Stocker, T.F., Qin, D., Plattner, G.K., Tignor, M., Allen, S.K., Boschung, J., Nauels, A., Midgley, P.M., Bex, V. and Xia, Y. 2013 IPCC, 2013: Climate Change 2013: The Physical Science Basis. Working Group I Contribution to the IPCC 5th Assessment Report, IPCC, Cambridge University Press, Cambridge, United Kingdom and New York, NY, USA, 1535 pp.

Streese, J. and Stegmann, R. 2003. Microbial oxidation of methane from old landfills in biofilters. Waste Manage 23(7), 573–580.

Svenning, M.M., Hestnes, A.G., Wartiainen, I., Stein, L.Y., Klotz, M.G., Kalyuzhnaya, M.G., Spang, A., Bringel, F., Vuilleumier, S., Lajus, A., Medigue, C., Bruce, D.C., Cheng, J.F., Goodwin, L., Ivanova, N., Han, J., Han, C.S., Hauser, L., Held, B., Land, M.L., Lapidus, A., Lucas, S., Nolan, M., Pitluck, S. and Woyke, T. 2011. Genome Sequence of the Arctic Methanotroph Methylobacter tundripaludum SV96. J Bacteriol 193(22), 6418–6419.

Tatari, K., Musovic, S., Gulay, A., Dechesne, A., Albrechtsen, H.J. and Smets, B.F. 2017. Density and distribution of nitrifying guilds in rapid sand filters for drinking water production: Dominance of Nitrospira spp. Water Res 127, 239–248.

Tatusov, R.L., Koonin, E.V. and Lipman, D.J. 1997. A genomic perspective on protein families. Science 278(5338), 631–637.

Tavormina, P.L., Orphan, V.J., Kalyuzhnaya, M.G., Jetten, M.S.M. and Klotz, M.G. 2011. A novel family of functional operons encoding methane/ammonia monooxygenase-related proteins in gammaproteobacterial methanotrophs. Environ Microbiol Rep 3(1), 91–100.

Tekerlekopoulou, A.G., Vasiliadou, I.A. and Vayenas, D.V. 2006. Physico-chemical and biological iron removal from potable water. Biochem Eng J 31(1), 74–83.

Trifinopoulos, J., Nguyen, L.T., von Haeseler, A. and Minh, B.Q. 2016. W-IQ-TREE: a fast online phylogenetic tool for maximum likelihood analysis. Nucleic Acids Res 44(W1), W232–W235.

Trussell, R.R., Hand, D.W., Tchobanoglous, G., Howe, K.J. and Crittenden, J.C. (2012) Principles of Water Treatment, John Wiley & Sons Inc, Hoboken, New Jersey.

van der Wielen, P.W.J.J., Voost, S. and van der Kooij, D. 2009. Ammonia-Oxidizing Bacteria and Archaea in Groundwater Treatment and Drinking Water Distribution Systems. Appl Environ Microbiol 75(14), 4687–4695.

Vaser, R., Sovic, I., Nagarajan, N. and Sikic, M. 2017. Fast and accurate de novo genome assembly from long uncorrected reads. Genome Res 27(5), 737–746.

Vekeman, B., Kerckhof, F.M., Cremers, G., de Vos, P., Vandamme, P., Boon, N., Op den Camp, H.J.M. and Heylen, K. 2016. New Methyloceanibacter diversity from North Sea sediments includes methanotroph containing solely the soluble methane monooxygenase. Environ Microbiol 18(12), 4523–4536.

Wagner, F.B., Nielsen, P.B., Boe-Hansen, R. and Albrechtsen, H.J. 2016. Copper deficiency can limit nitrification in biological rapid sand filters for drinking water production. Water Res 95, 280–288.

Wallar, B.J. and Lipscomb, J.D. 1996. Dioxygen activation by enzymes containing binuclear non-heme iron clusters. Chem Rev 96(7), 2625–2657.

Wang, Y., Ma, L., Mao, Y., Jiang, X., Xia, Y., Yu, K., Li, B. and Zhang, T. 2017. Comammox in drinking water systems. Water Res 116, 332–341.

Wang, Z.H., Cao, Y.Q., Zhu-Barker, X., Nicol, G.W., Wright, A.L., Jia, Z.J. and Jiang, X.J. 2019. Comammox Nitrospira clade B contributes to nitrification in soil. Soil Biol Biochem 135, 392–395.

Ward, N.L., Challacombe, J.F., Janssen, P.H., Henrissat, B., Coutinho, P.M., Wu, M., Xie, G., Haft, D.H., Sait, M., Badger, J., Barabote, R.D., Bradley, B., Brettin, T.S., Brinkac, L.M., Bruce, D., Creasy, T., Daugherty, S.C., Davidsen, T.M., Deboy, R.T., Detter, J.C., Dodson, R.J., Durkin, A.S., Ganapathy, A., Gwinn-Giglio, M., Han, C.S., Khouri, H., Kiss, H., Kothari, S.P., Madupu, R., Nelson, K.E., Nelson, W.C., Paulsen, I., Penn, K., Ren, Q.H., Rosovitz, M.J., Selengut, J.D., Shrivastava, S., Sullivan, S.A., Tapia, R., Thompson, L.S., Watkins, K.L., Yang, Q., Yu, C.H., Zafar, N., Zhou, L.W. and Kuske, C.R. 2009. Three Genomes from the Phylum Acidobacteria Provide Insight into the Lifestyles of These Microorganisms in Soils. Appl Environ Microbiol 75(7), 2046–2056.

Watanabe, T., Kojima, H., Umezawa, K., Hori, C., Takasuka, T.E., Kato, Y. and Fukui, M. 2019. Genomes of Neutrophilic Sulfur-Oxidizing Chemolithoautotrophs Representing 9 Proteobacterial Species From 8 Genera. Front Microbiol 10(316).

Wegner, C.E., Gaspar, M., Geesink, P., Herrmann, M., Marz, M. and Kusel, K. 2019. Biogeochemical Regimes in Shallow Aquifers Reflect the Metabolic Coupling of the Elements Nitrogen, Sulfur, and Carbon. Appl Environ Microbiol 85(9), 0099–2240.

Whitman, W.B., Oren, A., Chuvochina, M., da Costa, M.S., Garrity, G.M., Rainey, F.A., Rossello-Mora, R., Schink, B., Sutcliffe, I., Trujillo, M.E. and Ventura, S. 2018. Proposal of the suffix -ota to denote phyla. Addendum to ‘Proposal to include the rank of phylum in the International Code of Nomenclature of Prokaryotes’. Int. J. Syst. Evol. Microbiol. 68(3), 967–969.

Wickham, H. 2016 ggplot2: Elegant Graphics for Data Analysis, (Springer, Heidelberg), Springer, Heidelberg.

Wilczak, A., Jacangelo, J.G., Marcinko, J.P., Odell, L.H., Kirmeyer, G.J. and Wolfe, R.L. 1996. Occurrence of nitrification in chloraminated distribution systems. J Am Water Works Assoc 88(7), 74–85.

Yan, L., Herrmann, M., Kampe, B., Lehmann, R., Totsche, K.U. and Küsel, K. 2020. Environmental selection shapes the formation of near-surface groundwater microbiomes. Water Res 170, 115341.

Yang, J.X., Ma, J., Song, D., Zhai, X.D. and Kong, X.J. 2016. Impact of preozonation on the bioactivity and biodiversity of subsequent biofilters under low temperature conditions-A pilot study. Front Environ Sci Eng 10(4).

Zhang, Y.F., Yang, Y., Liu, J.S. and Qiu, G.Z. 2013. Isolation and characterization of Acidithiobacillus ferrooxidans strain QXS-1 capable of unusual ferrous iron and sulfur utilization. Hydrometallurgy 136, 51–57.

Zhou, J.Z., Bruns, M.A. and Tiedje, J.M. 1996. DNA recovery from soils of diverse composition. Appl Environ Microbiol 62(2), 316–322.

