## Supplementary material for "Metagenomic profiling of ammonia- and methane-oxidizing microorganisms in a Dutch drinking water treatment plant"

### Supplementary methods

#### *HMM profiles*

HMM profiles were generated by downloading protein sequences from the TrEMBL database of EMBL and linking them to KEGG (Ogata et al., 1999) numbers with the UniProt (Bateman et al., 2017) identifiers. For each K-number, the proteins were clustered using Linclust (Steinegger and Soding, 2018) (--kmer-per-seq 160 --min-seq-id 0.5 --similarity-type 1 --sub-mat blosum80 --cluster-mode 2 --cov-mode 0 -c 0.7). The proteins were aligned with MAFFT v7.397 (Katoh and Standley, 2013) (--anysymbol), and HMM profiles were created with hmmbuild tool within HMMER.

#### *16S rRNA analyses*

The 16S rRNA gene-based phylogenetic composition of the paired-end libraries were determined using phyloFlash v3.3b1 (Gruber-Vodicka, 2019) with SILVA_132_SSURef_NR99 as the reference database.

#### *Phylogenomic analyses*

Reference genomes (Table S2) (>90% completeness and <10% redundancy) were downloaded from NCBI and dereplicated at 99% ANI using dRep (Olm et al., 2017). All bacterial genome-based phylogenetic analyses were performed using the UBCG pipeline for extracting the core gene set for phylogenomic tree reconstruction (UBCG; (Na et al., 2018)). Extracted genes were aligned and concatenated using UBCG with default parameters. Maximum likelihood phylogenetic trees were calculated from the concatenated nucleotide alignment using RAxML version 8.2.10 (Stamatakis, 2014) on the CIPRES science gateway (Miller et al., 2010) with the GTR substitution and GAMMA rate heterogeneity models and 100 bootstrap iterations. For the archaeal phylogeny, the anvi'o phylogenomic workflow (http://merenlab.org/2017/06/07/phylogenomics) was used to individually align and concatenate 162 archaeal single-copy core genes (Rinke et al., 2013). The concatenated amino acid alignment was used to generate a phylogenetic tree by RAxML on CIPRES with the PROTGAMMA model and 100 bootstrap replications. All the phylogenetic trees were visualized in iTOL (Letunic and Bork, 2016).

*Phylogenetic analyses of key genes*

In total, 17 sequences classified as AmoA and 32 sequences classified as PmoA/PxmA with a length of ≥ 200 amino acids were included in the phylogenetic analysis. The sequences were imported into custom-made database and were aligned using the software package ARB v5.5 (Ludwig et al., 2004). For monooxygenases, a maximum likelihood tree including 286 aligned protein sequences was calculated using RaxML HPC-HYBRID v.8.2.12 on the CIPRES Science Gateway v.3.3 (Miller et al., 2010) with the LG substitution and GAMMA heterogeneity models and 1,000 bootstrap iterations. The resulting tree was visualized using the online tool iTOL (Letunic and Bork, 2016).

For nitrate reductases/nitrite oxidoreductases, all sequences classified as NarG or NxrA with a protein length ≥ 850 amino acid were used for phylogenetic analysis. In total, 36 sequences were imported into a custom-made database containing all groups of publicly available nitrate reductases and nitrite oxidoreductases. Sequences were aligned using the software package ARB v5.5 (Ludwig et al., 2004). A consensus maximum likelihood tree based on 375 aligned protein sequences was calculated using the IQ-tree webserver (Trifinopoulos et al., 2016) with the determined best-fit substitution model LG+I+G4 and 1,000 ultrafast bootstrap iterations. The constructed trees were imported into iTOL (Letunic and Bork, 2016) for visualization.

### Supplementary Figures

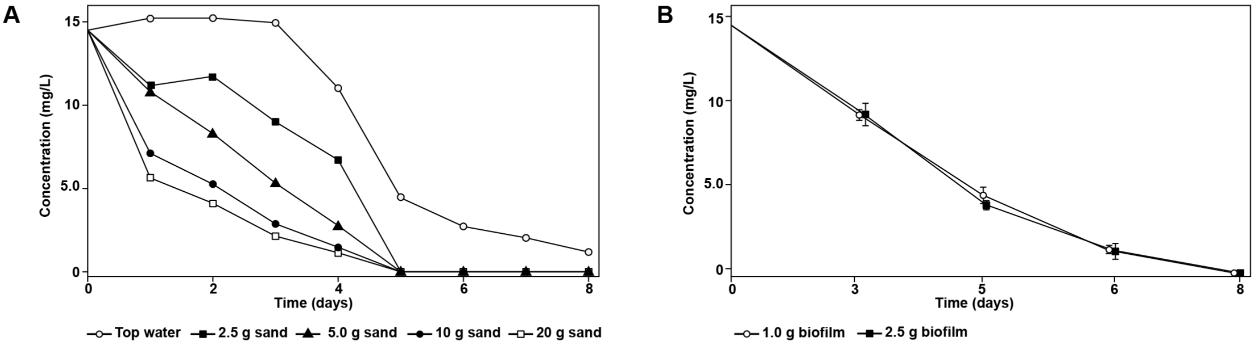


**Figure S1.** Methane oxidation activity in **(A)** P-RSF and **(B)** WB samples.

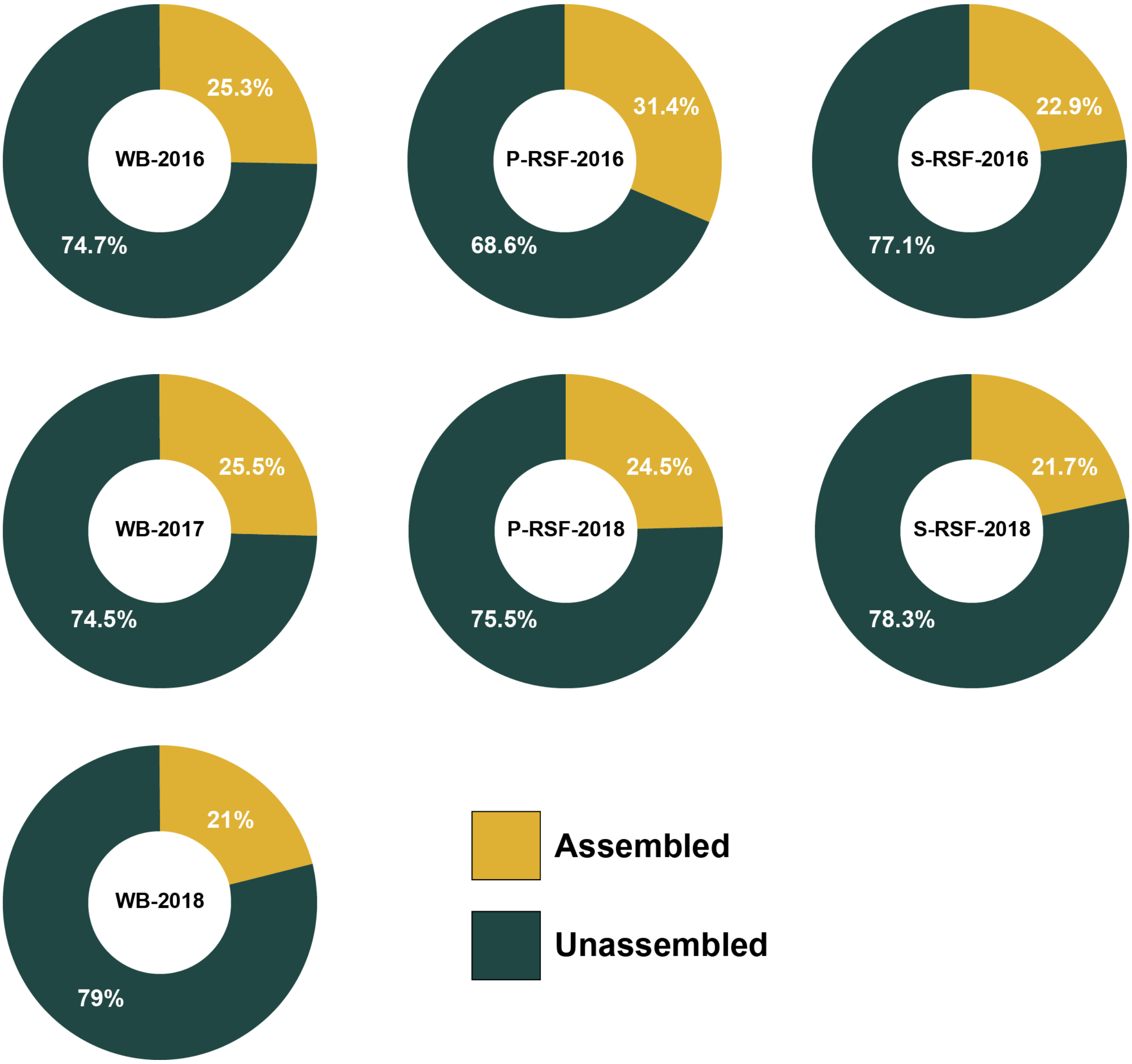


**Figure S2.** Donut charts illustrating the proportion of assembled and unassembled 16S rRNA reads in each sample.

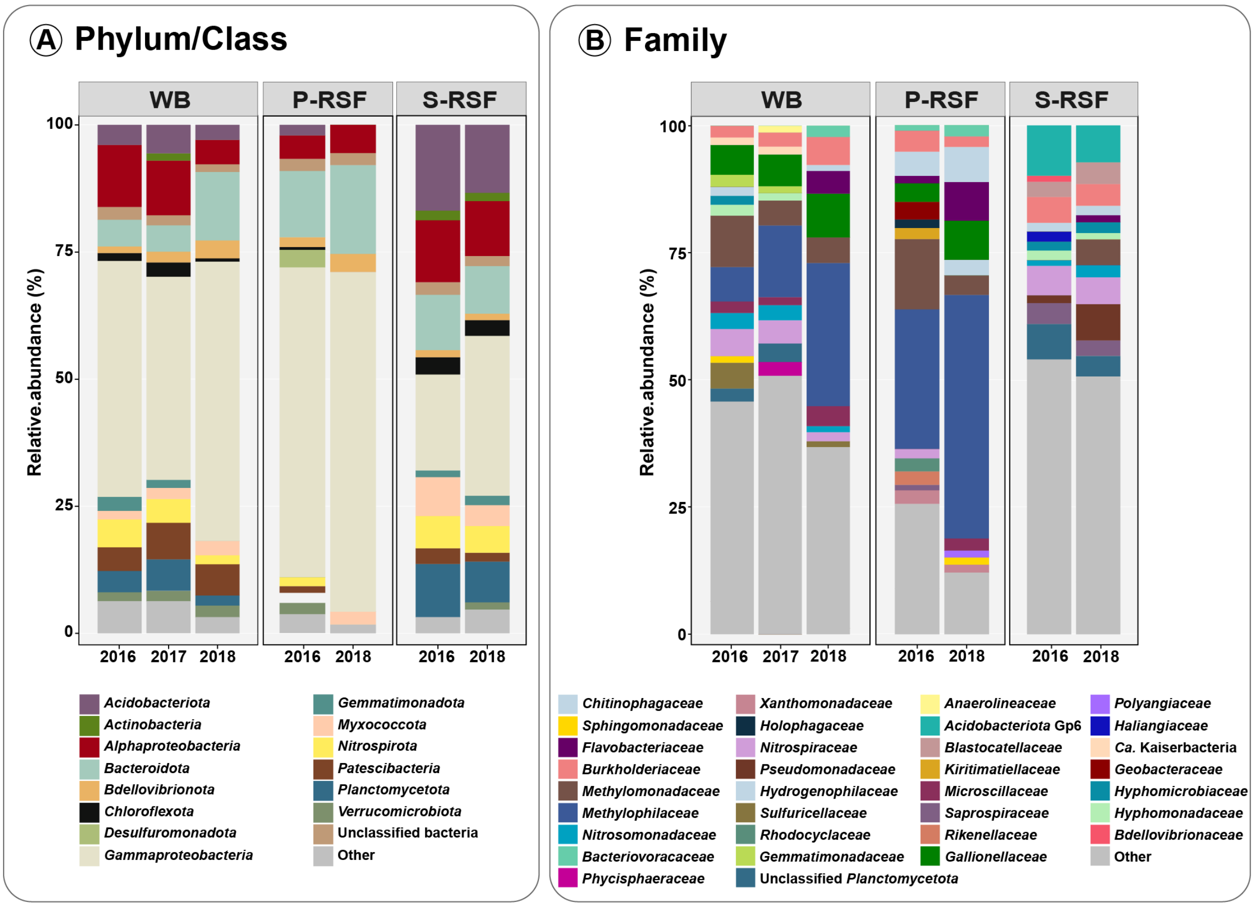


**Figure S3.** Bar charts illustrating the 16S rRNA gene-based relative abundance of bacterial taxa at **(A)** phylum and **(B)** family-level in each sample. Taxa represented by <1% of the total sequences were grouped as 'Other'.

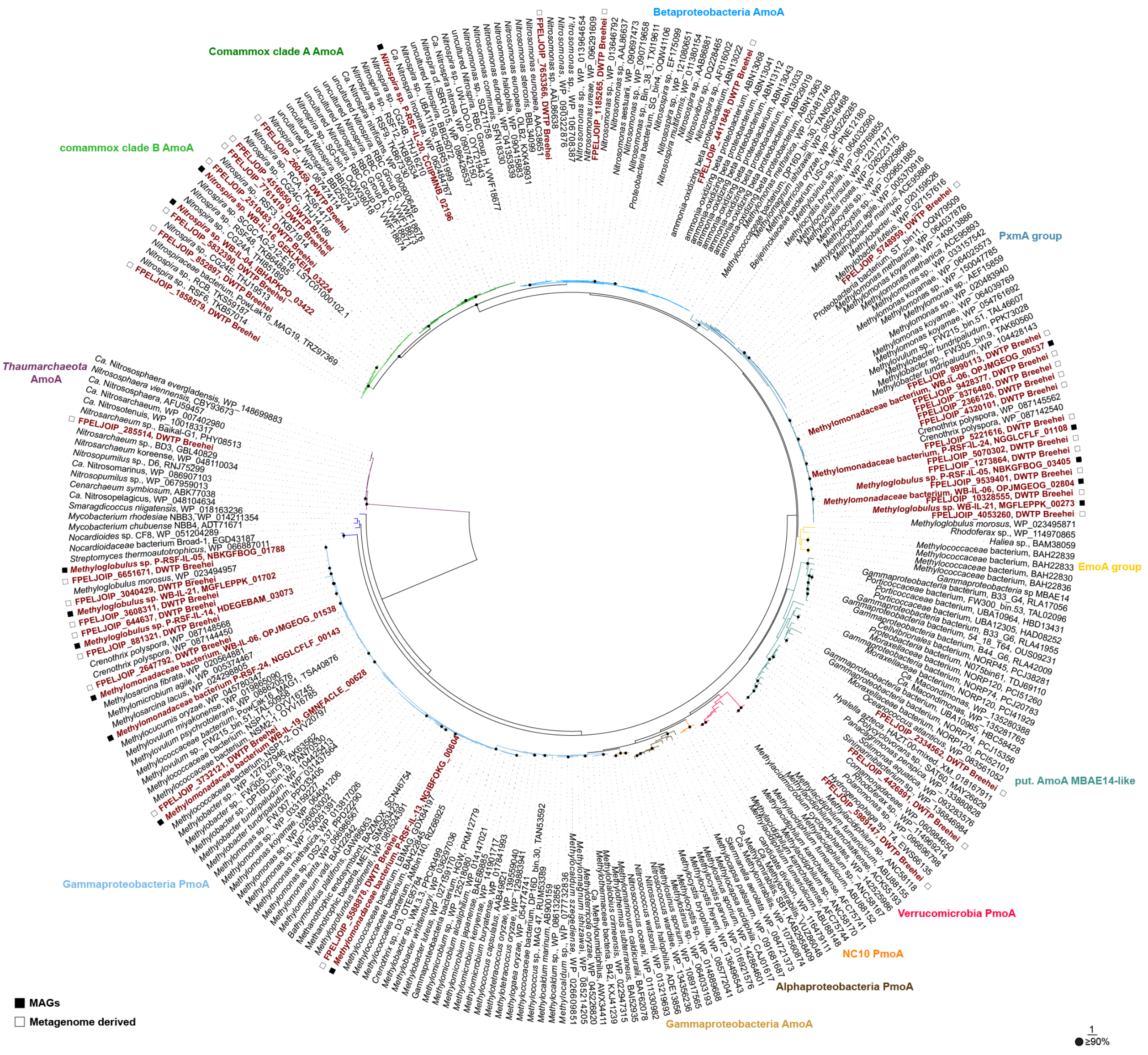


**Figure S4.** XmoA-based phylogenetic analysis of DWTP Breehei-derived and related protein sequences. Maximum likelihood phylogenetic tree was calculated based on 286 aligned XmoA sequences with a length ≥200 amino acids. Bootstrap support values ≥90% are indicated by black dots; the scale bar indicates estimated amino acid substitutions. Sequences obtained in this study are shown in red. Amo, ammonia monooxygenase; Emo, ethane monooxygenase; Pmo/Pxm, particulate methane monooxygenase.

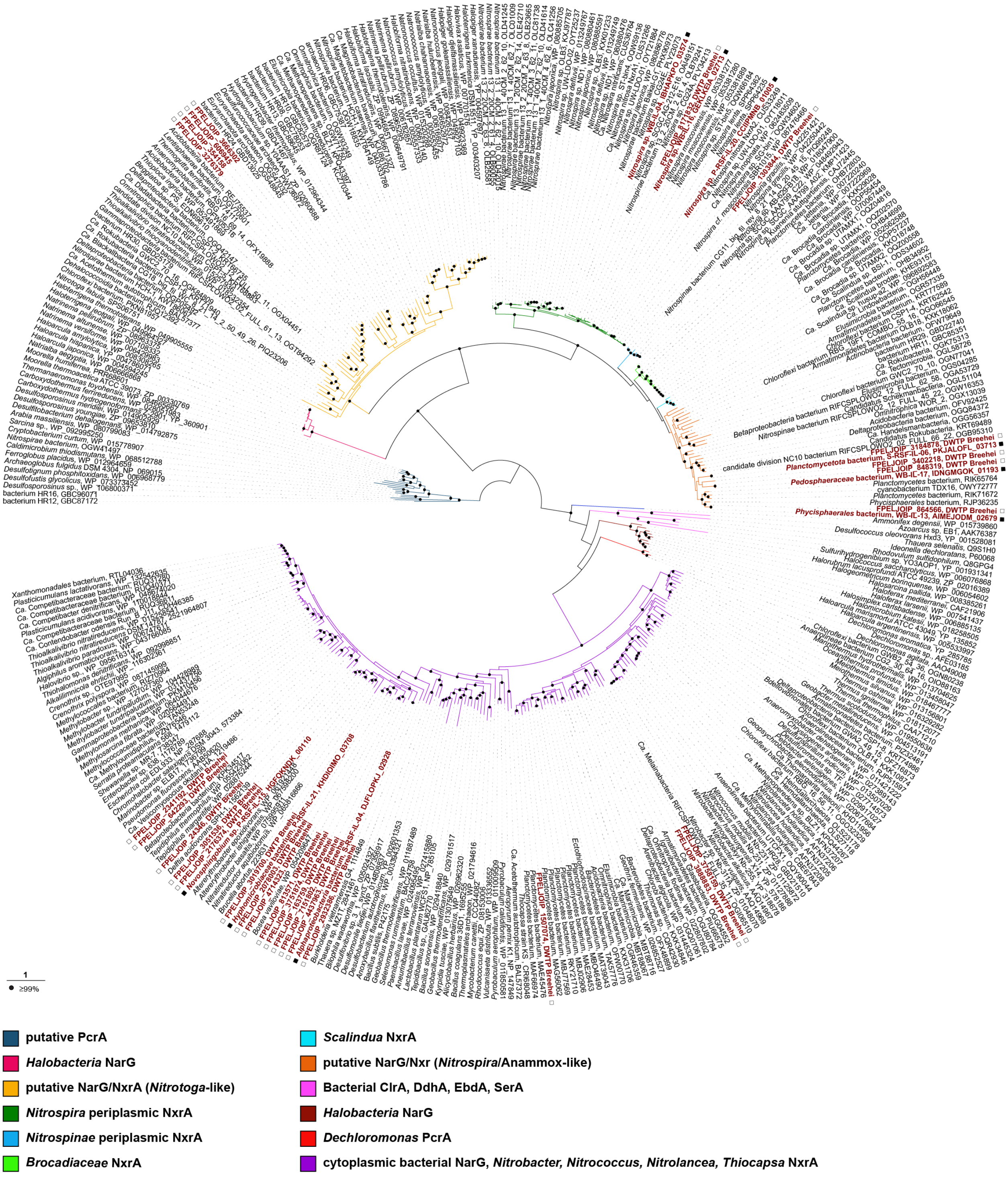


**Figure S5.** NxrA/NarG-based phylogenetic analysis of DWTP Breehei-derived and related protein sequences. Maximum likelihood phylogenetic tree of 424 bacterial NxrA and NarG sequences. Only sequences with a length ≥850 amino acid were included in the analysis. Bootstrap support values ≥99% are indicated by black dots; the scale bar indicates estimated amino acid substitutions. Sequences obtained in this study are shown in red. Clr, chlorate reductase; Ddh, dimethylsulfide dehydrogenase; Ebd, ethylbenzene dehydrogenase; Nar, nitrate reductases; Nxr, nitrite oxidoreductase; Pcr, perchlorate reductase; Ser, selenate reductase.

### Supplementary Tables

**Table S1.** DWTP Breehei metagenome assembly statistics.

| Statistics | WB | P-RSF | S-RSF |
|---|---|---|---|
| **Total number of reads** | 70,405,383 | 28,893,831 | 26,119,721 |
| **Number of assembled reads (%)[a]** | 56,675,821 (80.5%) | 25,060,071 (86.7%) | 18,960,046 (72.6%) |
| **Number of assembled reads assigned to MAGs (%)[b]** | 13,005,894 (23%) | 16,043,421 (64%) | 2,653,168 (14%) |
| **Total length (bp)** | 413,369,054 | 184,807,343 | 249,648,242 |
| **Number of contigs** | 132,794 | 41,634 | 81,371 |
| **Longest contig** | 396,269 | 630,225 | 317,318 |
| **Shortest contig** | 1,500 | 1,500 | 1,500 |
| **N50** | 3,101 | 5,132 | 3,112 |

[a]Based on bwa mapping (Li and Durbin, 2010)
[b]Number of reads assigned by CheckM coverage (Parks et al., 2015)

*Supplementary Tables supplied as separate Excel sheets*

**Table S2.** General overview of MAGs reconstructed from DWTP Breehei metagenomes.

**Table S3.** Presence/absence of genes involved in nitrification, methane and C1 oxidation, and sulfur cycling. ALDH, aldehyde dehydrogenase; AMO, ammonia monooxygenase (clade A and B comammox *Nitrospira*); FCC, flavocytochrome *c* sulfide dehydrogenase; FDH, formate dehydrogenase; MTHFD, methylenetetrahydrofolate dehydrogenase/cyclohydrolase; GSH, glutathione-linked formaldehyde oxidation; HAO, hydroxylamine dehydrogenase; HDR, heterodisulfide reductase; $H_4MPT$, tetrahydromethanopterin; HURM, hydroxylamineubiquinone reaction module; pMMO, particulate methane monooxygenase; sMMO, soluble methane monooxygenase; MDH, lanthanide and calcium-dependent methanol dehydrogenases; NAR, nitrate reductase; NXR, nitrite oxidoreductase; SOE, sulfite oxidase; SOX, sulfur oxidation system, SQR, sulfide:quinone oxidoreductase.

**Table S4.** List of ribosomal proteins used by the anvi′o phylogenomics workflow to calculate the FastTree for clustering the DWTP Breehei MAGs (Figure 3).

**Table S5.** List of reference genomes used for the phylogenomic analyses.

### References

Bateman, A., Martin, M.J., O'Donovan, C., Magrane, M., Alpi, E., Antunes, R., Bely, B., Bingley, M., Bonilla, C., Britto, R., Bursteinas, B., Bye-A-Jee, H., Cowley, A., Da Silva, A., De Giorgi, M., Dogan, T., Fazzini, F., Castro, L.G., Figueira, L., Garmiri, P., Georghiou, G., Gonzalez, D., Hatton-Ellis, E., Li, W.Z., Liu, W.D., Lopez, R., Luo, J., Lussi, Y., MacDougall, A., Nightingale, A., Palka, B., Pichler, K., Poggioli, D., Pundir, S., Pureza, L., Qi, G.Y., Rosanoff, S., Saidi, R., Sawford, T., Shypitsyna, A., Speretta, E., Turner, E., Tyagi, N., Volynkin, V., Wardell, T., Warner, K., Watkins, X., Zaru, R., Zellner, H., Xenarios, I., Bougueleret, L., Bridge, A., Poux, S., Redaschi, N., Aimo, L., Argoud-Puy, G., Auchincloss, A., Axelsen, K., Bansal, P., Baratin, D., Blatter, M.C., Boeckmann, B., Bolleman, J., Boutet, E., Breuza, L., Casal-Casas, C., de Castro, E., Coudert, E., Cuche, B., Doche, M., Dornevil, D., Duvaud, S., Estreicher, A., Famiglietti, L., Feuermann, M., Gasteiger, E., Gehant, S., Gerritsen, V., Gos, A., Gruaz-Gumowski, N., Hinz, U., Hulo, C., Jungo, F., Keller, G., Lara, V., Lemercier, P., Lieberherr, D., Lombardot, T., Martin, X., Masson, P., Morgat, A., Neto, T., Nouspikel, N., Paesano, S., Pedruzzi, I., Pilbout, S., Pozzato, M., Pruess, M., Rivoire, C., Roechert, B., Schneider, M., Sigrist, C., Sonesson, K., Staehli, S., Stutz, A., Sundaram, S., Tognolli, M., Verbregue, L., Veuthey, A.L., Wu, C.H., Arighi, C.N., Arminski, L., Chen, C.M., Chen, Y.X., Garavelli, J.S., Huang, H.Z., Laiho, K., McGarvey, P., Natale, D.A., Ross, K., Vinayaka, C.R., Wang, Q.H., Wang, Y.Q., Yeh, L.S., Zhang, J. and UniProt, C. 2017. UniProt: the universal protein knowledgebase. Nucleic Acids Res 45(D1), D158-D169.

Gruber-Vodicka, H.R., Seah, Brandon K. B., Pruesse, Elmar 2019 phyloFlash – Rapid SSU rRNA profiling and targeted assembly from metagenomes, p. 521922 bioRxiv.

Katoh, K. and Standley, D.M. 2013. MAFFT Multiple Sequence Alignment Software Version 7: Improvements in Performance and Usability. Mol Biol Evol 30(4), 772-780.

Letunic, I. and Bork, P. 2016. Interactive tree of life (iTOL) v3: an online tool for the display and annotation of phylogenetic and other trees. Nucleic Acids Research 44(W1), W242-W245.

Li, H. and Durbin, R. 2010. Fast and accurate long-read alignment with Burrows-Wheeler transform. Bioinformatics 26(5), 589-595.

Ludwig, W., Strunk, O., Westram, R., Richter, L., Meier, H., Yadhukumar, Buchner, A., Lai, T., Steppi, S., Jobb, G., Forster, W., Brettske, I., Gerber, S., Ginhart, A.W., Gross, O., Grumann, S., Hermann, S., Jost, R., Konig, A., Liss, T., Lussmann, R., May, M., Nonhoff, B., Reichel, B., Strehlow, R., Stamatakis, A., Stuckmann, N., Vilbig, A., Lenke, M., Ludwig, T., Bode, A. and Schleifer, K.H. 2004. ARB: a software environment for sequence data. Nucleic Acids Res 32(4), 1363-1371.

Miller, M.A., Pfeiffer, W. and Schwartz, T. 2010 Creating the CIPRES Science Gateway for inference of large phylogenetic trees. Gateway Computing Environments Workshop (GCE), pp. 1-8.

Na, S.I., Kim, Y.O., Yoon, S.H., Ha, S.M., Baek, I. and Chun, J. 2018. UBCG: Up-to-date bacterial core gene set and pipeline for phylogenomic tree reconstruction. J Microbiol 56(4), 280-285.

Ogata, H., Goto, S., Sato, K., Fujibuchi, W., Bono, H. and Kanehisa, M. 1999. KEGG: Kyoto Encyclopedia of Genes and Genomes. Nucleic Acids Res 27(1), 29-34.

Olm, M.R., Brown, C.T., Brooks, B. and Banfield, J.F. 2017. dRep: a tool for fast and accurate genomic comparisons that enables improved genome recovery from metagenomes through de-replication. ISME J 11(12), 2864-2868.

Parks, D.H., Imelfort, M., Skennerton, C.T., Hugenholtz, P. and Tyson, G.W. 2015. CheckM: assessing the quality of microbial genomes recovered from isolates, single cells, and metagenomes. Genome research 25(7), 1043-1055.

Rinke, C., Schwientek, P., Sczyrba, A., Ivanova, N.N., Anderson, I.J., Cheng, J.F., Darling, A., Malfatti, S., Swan, B.K., Gies, E.A., Dodsworth, J.A., Hedlund, B.P., Tsiamis, G., Sievert,

S.M., Liu, W.T., Eisen, J.A., Hallam, S.J., Kyrpides, N.C., Stepanauskas, R., Rubin, E.M., Hugenholtz, P. and Woyke, T. 2013. Insights into the phylogeny and coding potential of microbial dark matter. Nature 499(7459), 431-437.

Stamatakis, A. 2014. RAxML version 8: a tool for phylogenetic analysis and post-analysis of large phylogenies. Bioinformatics 30(9), 1312-1313.

Steinegger, M. and Soding, J. 2018. Clustering huge protein sequence sets in linear time. Nat Commun 9(2041-1723).

Trifinopoulos, J., Nguyen, L.T., von Haeseler, A. and Minh, B.Q. 2016. W-IQ-TREE: a fast online phylogenetic tool for maximum likelihood analysis. Nucleic Acids Res 44(W1), W232-W235.
